# Structural Repetition Detector: multi-scale quantitative mapping of molecular complexes through microscopy

**DOI:** 10.1101/2024.09.16.613204

**Authors:** Afonso Mendes, Bruno M. Saraiva, Guillaume Jacquemet, João I. Mamede, Christophe Leterrier, Ricardo Henriques

## Abstract

From molecules to organelles, cells exhibit recurring structural motifs across multiple scales. Understanding these structures provides insights into their functional roles. While super-resolution microscopy can visualise such patterns, manual detection in large datasets is challenging and biased. We present the Structural Repetition Detector (SReD), an unsupervised computational framework that identifies repetitive biological structures by exploiting local texture repetition. SReD formulates structure detection as a similarity-matching problem between local image regions. It detects recurring patterns without prior knowledge or constraints on the imaging modality. We demonstrate SReD’s capabilities on various fluorescence microscopy images. Quantitative analyses of three datasets highlight SReD’s utility: estimating the periodicity of spectrin rings in neurons, detecting HIV-1 viral assembly, and evaluating microtubule dynamics modulated by EB3. Our open-source ImageJ and Fiji plugin enables unbiased analysis of repetitive structures across imaging modalities in diverse biological contexts.

## Introduction

Biological systems exhibit structural repetition across multiple scales, from biomolecules to supramolecular assemblies and cellular structures (1). Understanding these patterns is crucial for identifying their functional significance and underlying biological processes (2). Microscopy techniques offer molecular-level resolution but manually detecting repetitive motifs in large datasets is impractical, biased, and expertise-dependent (3). To address these limitations, machine learning, particularly deep convolutional neural networks (CNNs), has been employed to detect and segment biological structures automatically (4, 5). However, CNNs require extensive labelled training data, inheriting biases (6). Previous methods enable unbiased registration but need single-molecule localisation data, limiting their applicability (7, 8). We present the Structural Repetition Detector (SReD), an unsupervised framework to identify repetitive biological structures by exploring local texture redundancy. SReD formulates structure detection as similarity matching between local image regions, allowing pattern detection without prior knowledge or microscopy modality constraints. We demonstrate SReD’s capabilities on fluorescence microscopy images of diverse cell types and structures, including microtubule networks, nuclear envelope, pores, and virus particles (**Fig. 1**). SReD generates Structural Repetition Scores (SRSs) highlighting regions with repetitive textures. Users can provide artificial blocks or extract them from the data for repetition analysis. An unbiased sampling scheme maps global repetition by testing every possible image block as a reference (**Note S1**). We showcase SReD’s utility through three datasets: 1) spectrin rings in neuronal axons, accurately estimating ring periodicity and pinpointing periodic patterns, 2) HIV-1 Gag assembly, mapping viral structures without structural priors, and 3) dynamic EB3 and microtubule structures, assessing structural displacement and stability over time. Our open-source ImageJ and Fiji plugin enables versatile, unbiased analysis of redundancy in microscopy images. SReD advances computational microscopy by providing a generalised framework for detecting repetitive structures without labelled training data or single-molecule localisation input, facilitating the quantitative study of structural motifs across scales in diverse imaging datasets.

**Fig. 1.**
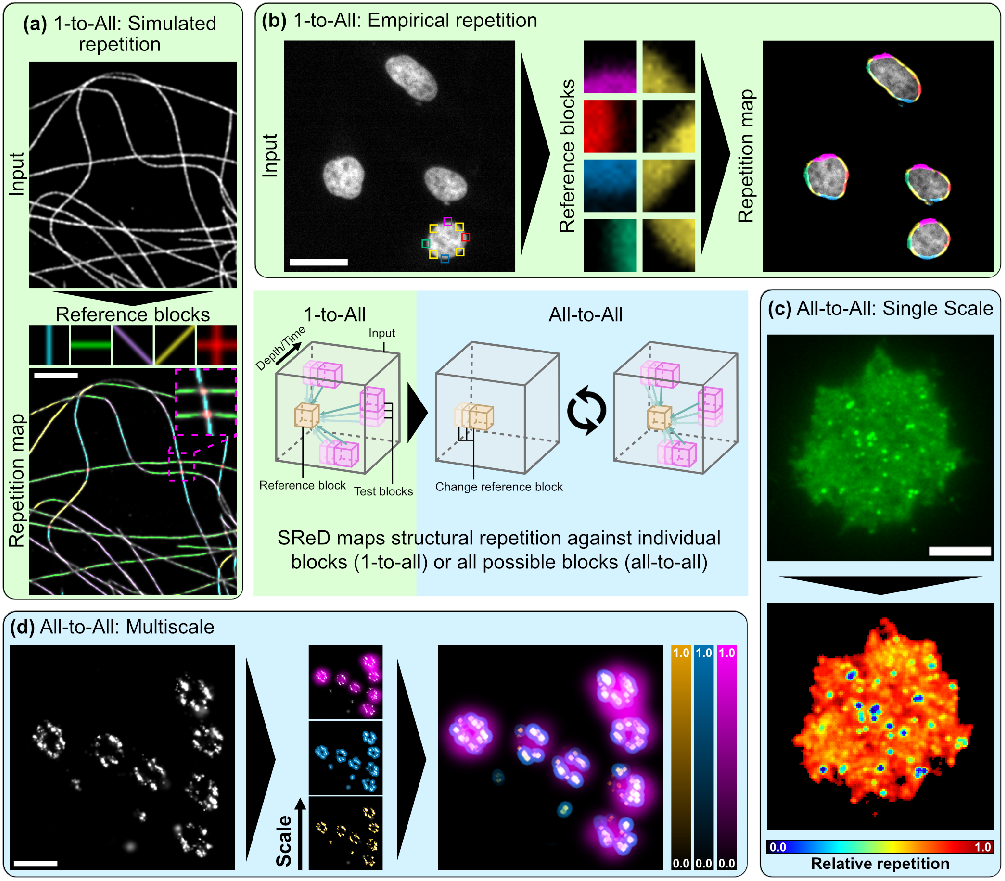
Applications of the Structural Repetition Detector (SReD) Algorithm in Fluorescence Microscopy. **a**, Detection of Structural Repetition Using Simulated Blocks: Microtubules imaged with STORM analysed for repetitive patterns using simulated structural blocks. Coloured regions in repetition map correspond to repetitions of same-coloured blocks above. **b**, Detection of Structural Repetition Using Empirical Blocks: HeLa cell nuclei stained with DAPI used to detect repetitive structural patterns using manually extracted empirical blocks. Coloured regions in repetition map correspond to repetitions of same-coloured blocks in previous subpanel. **c**, Global Repetition Detection: Jurkat cell expressing inducible HIV-1 Gag-EGFP fusion protein analysed using global repetition detection. Image probed for structural repetition using all possible empirical patches. Repetition map reveals structures not easily detectable in input image and their relative frequency. **d**, Multiscale Global Repetition: *Xenopus laevis* nuclear pores imaged with STORM analysed using different-sized receptive fields to detect structural repetition at various scales. Repetition map identifies repeated structures from single nucleoporins (orange) to nucleoporin clusters (blue) and nuclear pore units (magenta). Centre panel: Simplified SReD algorithm workflow, illustrating key steps from input preprocessing to repetition map generation.

## Results

### General applications of SReD

SReD is an open-source ImageJ and Fiji plugin that leverages GPU acceleration to identify repetitive patterns in microscopy images. The algorithm’s workflow, outlined in **Fig. 1** (centre panel) and **Note S1**, begins with the application of the Generalised Anscombe Transform (GAT) to stabilise noise variance (9). This step addresses the noise in microscopy images, which often exhibit Poisson and Gaussian noise. The GAT nonlinearly remaps pixel values to produce an image with near-Gaussian noise and stabilised variance, preserving local contrast and overall image statistics. This stabilisation is essential for robust downstream processing, mitigating violations of normality, homoscedasticity, and outlier assumptions that can compromise correlation metrics. Following noise stabilisation, SReD generates a relevance mask to exclude regions lacking substantive structural information (**Note S1**; **Fig. S1**). The analysis proceeds using reference blocks, either simulated or empirically sampled from the image. These blocks are matched against the input using correlation metrics to generate repetition maps, as detailed in the Methods section. The resulting repetition map highlights regions likely to contain structural repetitions, with nonlinear mapping applied to visually emphasise salient features. To demonstrate SReD’s versatility across diverse biological contexts, we conducted a comprehensive analysis of various microscopy datasets (**Fig. 1**; **Note S2**). We first examined a STORM image reconstruction of a cell with labelled microtubules (10). This approach effectively mapped microtubules at various orientations and crossings (**Fig. 1a**; **Fig. S2**). We further illustrate SReD’s versatility by detecting nuclear envelopes in DAPI-stained cells (11) using empirical reference blocks extracted directly from the input image, distinguishing different morphological states potentially related to cell division or stress (**Fig. 1b**; **Fig. S3**). SReD also enables characterisation of structures without user-provided references. We exemplify this functionality by analysing an image of a Jurkat cell expressing an HIV-1 Gag-EGFP construct, which induces the production of virus-like particles (VLPs) (**Fig. 1c**; **Fig. S4**). In this mode, SReD mapped every structure in the image and assigned scores based on their relative repetition. As expected, the top score was given to the most repeated element, the diffuse eGFP signal, with viral structures exhibiting lower frequencies. Localisation of round viral structures via local extrema calculation revealed that the repetition map provided a superior platform for extrema detection compared to direct analysis of the raw images. The algorithm’s multiscale analysis capability is achieved by adjusting the ratio of block-to-image dimensions. Larger ratios capture larger structures, while smaller ratios capture finer details. For computational efficiency, it is preferable to modulate scale by downscaling the input rather than enlarging blocks, although combining both approaches often preserves structural detail best. We demonstrate this multiscale analysis by examining nuclear pore complexes in STORM image reconstructions with labelled gp210 proteins (12). SReD successfully mapped structures across different scales, discerning single nucleoporins, nucleoporin clusters, and entire nuclear pores (**Fig. 1d**; **Fig. S5**).

### 1-to-all case example: detection of spectrin ring periodicity in axons

We used SReD’s block repetition mode to map and quantify the membrane-associated periodic scaffold (MPS) architecture in neuronal axons automatically and without bias (**Fig. 2**). The MPS, composed of actin, spectrin, and associated proteins, forms a crucial structural component of neuronal axons (13, 14). Super-resolution microscopy has shown that the MPS consists of ring-like structures spaced 180-190 nm apart, with alternating actin/adducin and spectrin rings orthogonal to the axon’s long axis (15). Mapping this nanoscale organisation across entire neuron samples has been challenging due to the need for manual region selection, potentially introducing bias. We analysed datasets from Vassilopoulos *et al*. (16), comparing neurons treated with DMSO (control) or swinholide A (SWIN, an actin-disrupting drug). Using SReD, we developed an automated workflow to determine axon orientations by probing skeletonised neuron images with simulated lines at varying angles (**Note S3**; **Fig. S6**). This enabled consistent alignment of axon segments for downstream analysis. We optimised parameters for simulated ring blocks to match observed ring patterns in control data, yielding an inter-ring spacing of 192 nm, consistent with previous studies (**Fig. S7**)(15, 16). SReD generated repetition maps highlighting regions of high local similarity across neuron samples, allowing automatic extraction and quantification of MPS organisation without manual region selection (**Fig. 2b**; **Fig. S8**). We measured an average spacing of 178 nm under control conditions (**Fig. 2c**; **Fig. S9d**). In agreement with Vassilopoulos et al. (16), repetition maps showed that swinholide A treatment disrupted MPS structure, with reduced pattern prominence and frequency compared to controls (**Fig. 2b,c**). We used correlation metrics that minimise information loss while being aware of potential imprinting. Nonlinear mapping effectively distinguishes real patterns from imprinted ones (**Fig. S2d,e**; **Fig. S9c**). Our method accounts for neuron thickness variability and provides the average distance between patterns for additional biological insights. SReD’s local repetition scores quantified the fraction of structures with MPS patterns, revealing a 39% reduction in axons with detectable periodic scaffolds after swinholide A treatment (P<0.001, **Fig. 2d**). SReD’s maps identified drug-affected regions with confidence values, offering a detailed platform for analysing structural dysregulation (**Fig. 2c**). SReD also showed higher statistical sensitivity, detecting a 12% reduction in pattern prominence post-treatment (P<0.05) previously unreported (**Fig. S9e**). To test SReD’s noise robustness, we conducted a sensitivity analysis with images at varying signal-to-noise ratios (SNRs) (**Fig. S10**). SReD consistently detected ring structures even at low SNRs near 1, where patterns were visually indiscernible. SReD-generated maps outperformed direct STORM reconstructions in autocorrelation analysis, reliably identifying an average inter-ring spacing of 192 nm across all SNRs, demonstrating the algorithm’s robustness in detecting structural periodicity despite significant noise. We assessed SReD’s specificity and robustness to pattern deformations by applying stretch deformations to test images (**Fig. S11**). As the stretch factor increased, the average SRS decreased, indicating pattern disruption. However, SReD remained specific to the original pattern within the expected interval. Even at higher stretch factors, non-specific patterns were quantitatively discernible and reflected the intrinsic properties of the test data. This robustness is valuable for analysing periodic structures in diverse biological contexts, where deviations from ideal patterns are common due to sample preparation artefacts, imaging noise, or biological variability.

**Fig. 2.**
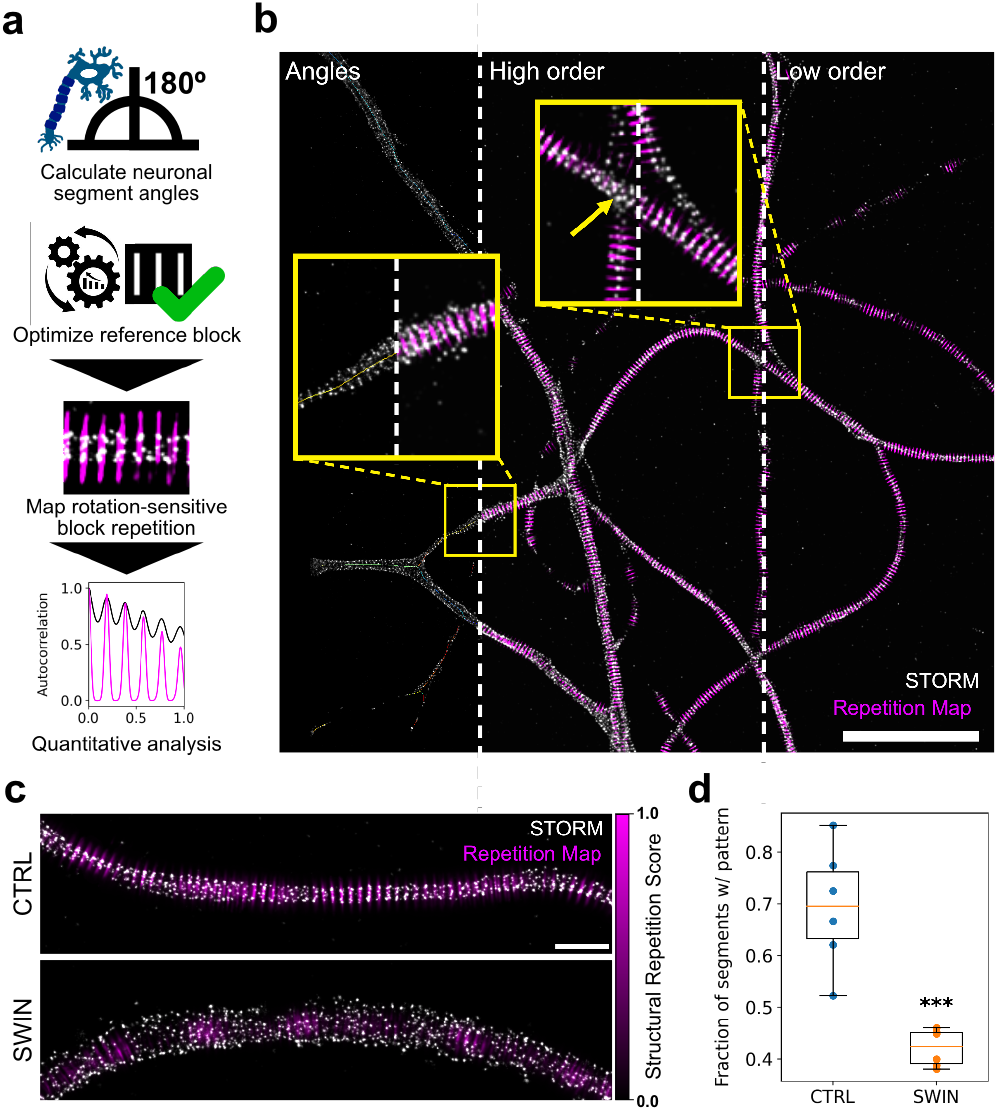
Automated detection and quantification of spectrin ring periodicity in neuronal axons. **a**, SReD-based analysis pipeline: the algorithm determines axon orientations, optimises a reference block for spectrin rings, and maps structural repetitions. Quantitative analysis is performed using autocorrelation and other methods. **b**, Control dataset image: STORM localization density (grey) overlaid with SReD repetition map (magenta). Insets: (i) ‘Angles’ -axon skeletons colour-coded by orientation; (ii) ‘High order’ - repetition map with a 9-ring reference block; (iii) ‘Low order’ - repetition map with a 3-ring reference block. Scale bar: 5 µm. **c**, Repetition maps comparing control (CTRL) and swinholide A-treated (SWIN) groups. SWIN-treated samples show reduced periodic structures. Scale bar: 1 µm. **d**, Quantification of axon segments with ring patterns. Bar graph shows a significant reduction in pattern-containing segments in SWIN vs. CTRL (n=6 per group, mean±SEM; CTRL: 0.694±0.008, SWIN: 0.421±0.007; p<0.001, unpaired t-test).

### All-to-all 3D case example: detecting HIV-1 Gag assembly in 3D

The establishment of a viral infection is the product of complex host-pathogen interactions, comprising an evolutionary “tug-of-war” where cells evolve protective mechanisms whilst viruses evolve to circumvent them. Viruses typically hijack cellular transcription and translation machinery to produce viral progeny required for viral replication (17). Therefore, viral assembly represents a critical platform for host-pathogen interactions that significantly impact infection outcomes. The HIV-1 *gag* gene encodes the Gag polypeptide precursor, which is cleaved into several key structural components. This polypeptide aggregates at the membrane of infected cells and induces the budding of mem-branous viral particles (17). Expression of Gag alone is sufficient to induce the formation of non-infectious virus-like particles (VLPs)(18, 19). To map viral structures in an unbiased manner, we examined an image of a Jurkat cell expressing an inducible HIV-1 Gag-EGFP construct using SReD (**Fig. 3a,b**). To evaluate the algorithm’s accuracy, we generated a population of simulated diffraction-limited particles with randomly distributed intensities across the image’s dynamic range, which served as a reference for comparison. Local maxima corresponding to active viral assembly sites were calculated from both the input image and the repetition map using identical parameters (**Fig. 3c**). Remarkably, SReD enabled the detection of 96% of the simulated particles, compared to only 32% in the input image, demonstrating the algorithm’s superior accuracy over direct analysis of input images (**Fig. 3d**). Visual inspection of the detected EGFP intensity signal vs. the SRS for the same pixel location revealed that high SRS regions corresponded to input regions with a wide range of intensity values. We observed that most structures of interest were allocated to the sample fraction above an arbitrary threshold of SRS 0.8, whilst the fraction below this threshold contained mostly background signal and some reference particles (**Fig. 3e**). Given that autofluorescence often corrupts microscopy analyses, we evaluated the algorithm’s performance in the presence of synthetic non-specific structures (**Note S4**). The repetition maps produced by SReD consistently provided a superior platform for detecting simulated reference particles and viral structures across conditions (**Fig. S12**). This analysis demonstrates SReD’s robust capability to map biological structures, such as assembling viral particles. The algorithm’s high sensitivity and specificity, even in the presence of non-specific structures, highlight its potential for studying dynamic cellular processes like viral assembly, where the ability to accurately detect and characterise structures amidst variable backgrounds is showcased.

**Fig. 3.**
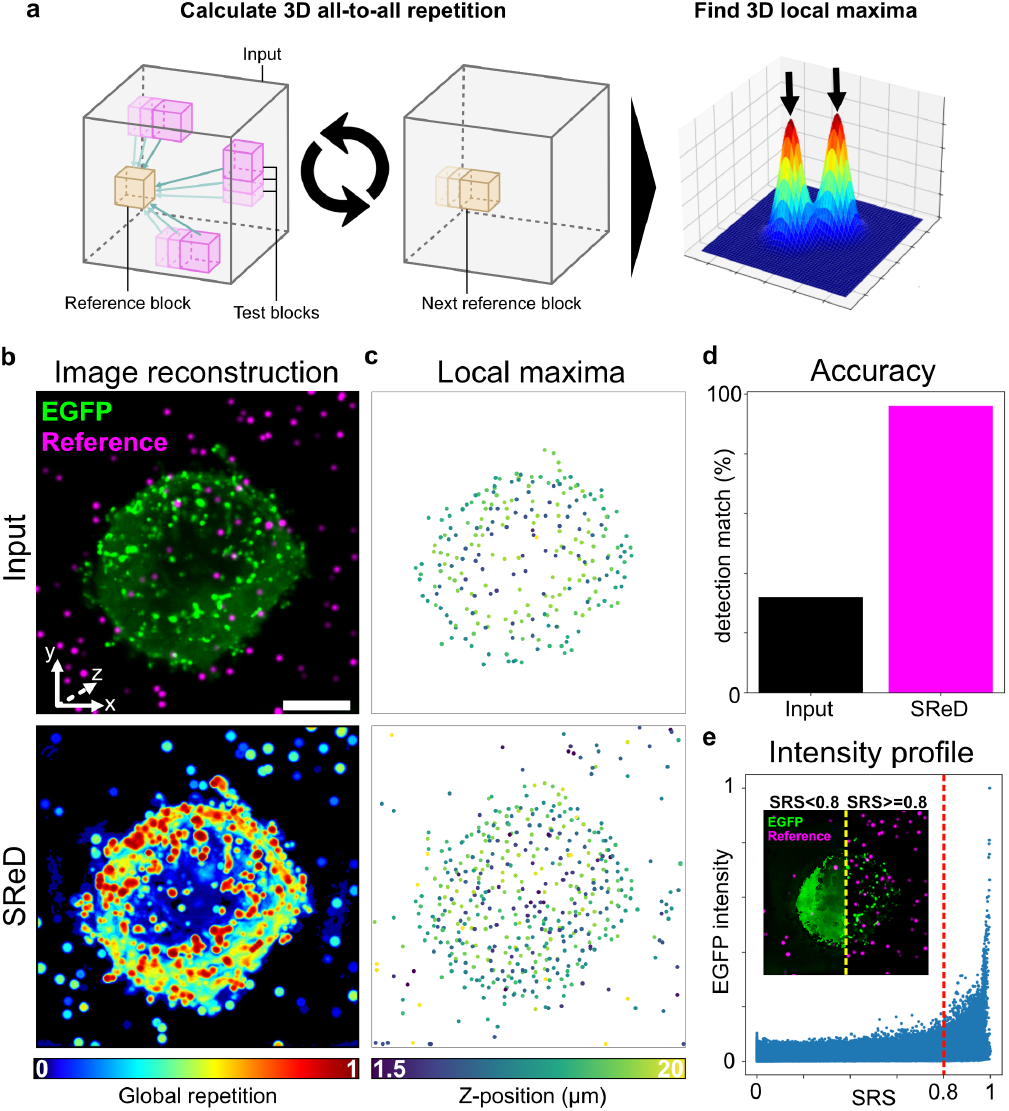
Detecting HIV-1 Virus-Like Particles in 3D. **a**, Analysis pipeline schematic. The algorithm uses 3D reference blocks for structural repetition analysis, locating viral structures via local maxima from the input image and repetition map. **b**, Z-projections of the input image (left) and repetition map (right), highlighting viral-like particles (“EGFP”) and simulated reference particles (“Reference”). **c**, Local maxima plots showing detected structures in the input image (left) and repetition map (right), with increased sensitivity in the repetition map. **d**, Accuracy plot comparing artificially added reference particle detection: input image (32%) vs. repetition map (96%). **e**, Intensity profile graph of EGFP signal (green) and structural repetition score (SRS, magenta), with a threshold at SRS 0.8 (dashed red line). Inset shows pixels below (dark) and above (light) the threshold, indicating high-SRS structures. Scale bars: 5 µm (main images), 1 µm (insets).

### All-to-all live-cell case example: assessment of microtubule dynamics

The multidimensional capabilities of SReD can be extended to analyse structural dynamics over time, providing insights into structural stability. We demonstrate this application using time-lapse imaging of RPE1 cells stably expressing End-binding Protein 3 (EB3) fused to GFP (**Fig. 4a**). EB3 binds to the plus ends of microtubules, appearing as comet-like structures that travel along the cytoplasm when visualised under fluorescence microscopy (20). We generated a global repetition map by treating time as the third dimension in our analysis, using a time-lapse sequence of approximately 2 minutes (**Fig. 4b**). To quantify structural changes, we calculated the Normalised Root Mean Squared Error (NRMSE) between the first and last frames of the time-lapse for both the input images and the repetition maps. The NRMSE of the input images reflected the spatial displacement of dynamic structures, yielding a relatively low value. In contrast, the NRMSE calculated from the repetition maps was substantially higher, indicating greater sensitivity to structural changes over time (**Fig. 4c**). SReD effectively mapped the spatial distribution of EB3 comet activity over time. By quantifying the repetitiveness of structures, it assigned scores to different regions, highlighting areas with high EB3 comet presence and their trajectories. The NRMSE maps further emphasised this distinction, revealing elevated values along comet paths, indicative of their dynamic nature. In contrast, the MTOC demonstrated notably lower NRMSE, suggesting its greater stability compared to the more mobile EB3 comets (**Fig. 4d**). The time interval used in the analysis captures the relatively slower dynamics of EB3 comets in this context. While individual comet tracking is not the primary focus of this method, the approach effectively reveals the spatiotemporal stability of structures, where instability often results from displacement, visually manifesting as comet trajectories. To further validate our approach, we performed the analysis with increased temporal resolution. We compared SReD’s results with conventional time projections of the input data, revealing advantages of our method. Unlike time projections, which typically integrate local intensities across time, SReD calculates local correlations of images across time, providing relative repetition scores that indicate how much the texture at each location changes relative to all other textures. This approach offers two significant benefits: (i) it provides a more nuanced measure of structural stability over time, and (ii) it is less susceptible to noise and intensity inconsistencies across time points (**Note S5; Fig. S13**). In this type of combined spatial and temporal analysis, instead of producing a time series, SReD’s output is a single map that shows the local stability of the timelapse over a specific time interval. This representation offers a comprehensive view of the structural dynamics that is not easily achieved using traditional methods such as kymographs. While kymographs are useful for tracking individual structures over time, SReD provides a broader perspective on the overall stability and dynamics of subcellular structures across the entire field of view.

**Fig. 4.**
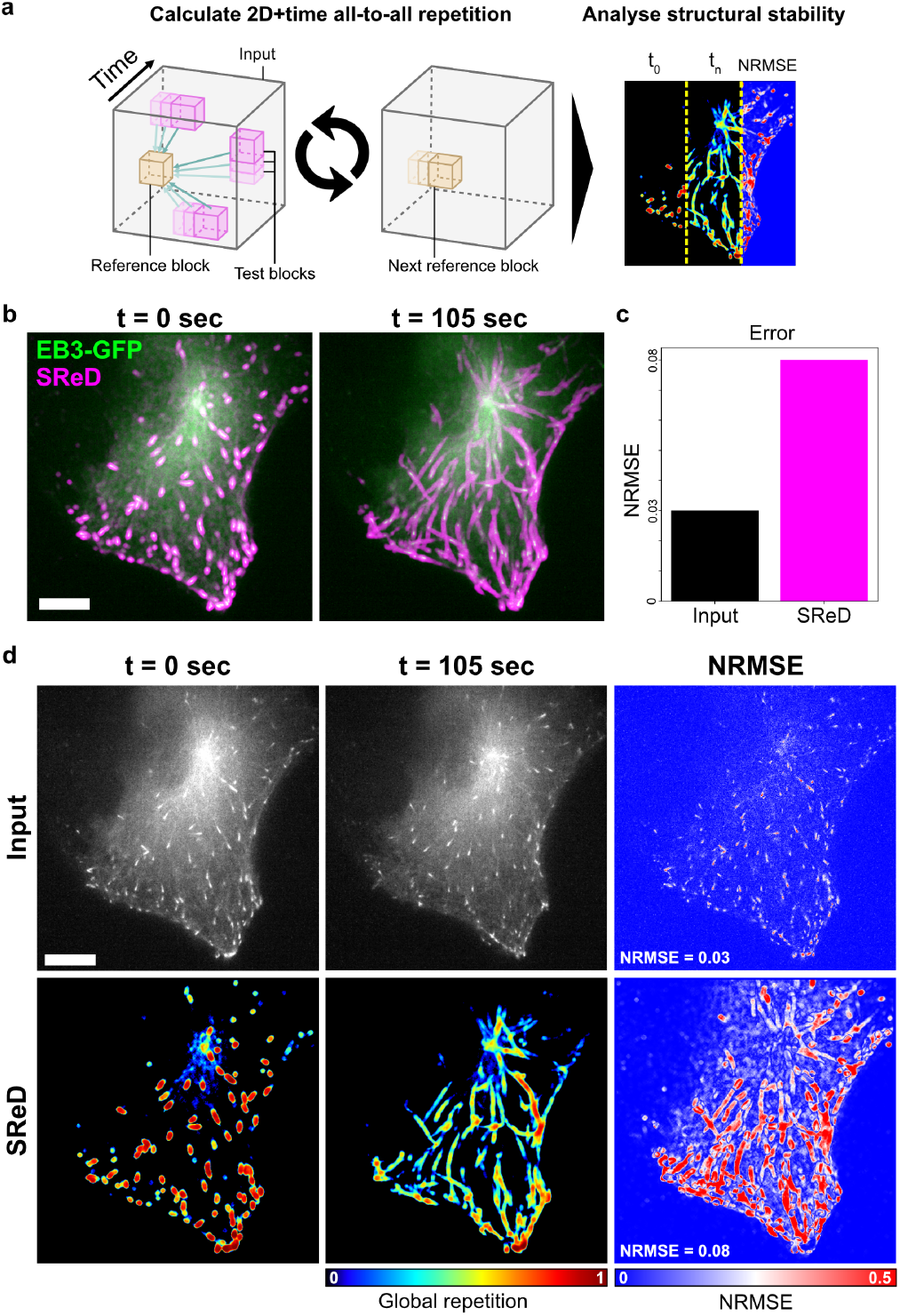
Assessment of microtubule dynamics using SReD. **a**, Analysis pipeline schematic. Global repetition analysis used time as the third dimension on a timelapse sequence of RPE1 cells expressing EB3-GFP over 105 seconds (35 frames). The first frame served as the control. Normalised Root Mean Squared Error (NRMSE) quantified structural differences between time points. **b**, Input images (left) and global repetition maps (right). The first frame’s repetition map highlights EB3 comets, while the entire time-lapse map shows comet trajectories and repetition over time. **c**, Bar graph of average NRMSE between input images and repetition maps. Higher error in repetition maps (0.08) vs. control images (0.03) indicates greater sensitivity to structural changes. **d**, NRMSE maps of input images (left) and repetition maps (right), showing structural stability over time. High NRMSE values (warmer colours) in EB3 trajectories indicate lower stability, while lower values (cooler colours) in the Microtubule Organising Centre (MTOC) indicate higher stability. Scale bars: 10 µm (main images), 2 µm (insets).

## Discussion and Conclusions

Our results demonstrate SReD’s versatility and analytical power across diverse biological contexts. In neuronal axons, SReD enabled automated, unbiased mapping of the membrane-associated periodic scaffold (MPS), revealing nuanced changes in pattern frequency and prominence following pharmacological perturbation. While previous studies by Vassilopoulos et al. (16) reported a 40% reduction in overall MPS prominence after treatment with swinholide A, SReD’s analysis provides a more detailed characterisation of the phenotype. By first distinguishing between regions with and without periodic patterns, and then analysing only the pattern-present areas, SReD detected a 12% reduction in pattern prominence and a 32% reduction in pattern frequency. This refined analysis not only corroborates the previously reported overall effect but also decomposes it into two distinct components, offering deeper insights into the nature of the structural changes. For HIV-1 Gag assembly, SReD achieved highly sensitive detection without relying on structural priors, significantly outperforming direct analysis of input images. This capability is particularly valuable in studying dynamic cellular processes like viral assembly, where the ability to accurately detect and characterise structures amidst variable backgrounds is crucial. In live-cell imaging of microtubule dynamics, SReD’s multidimensional capabilities allowed for quantitative assessment of structural stability across space and time. This analysis provided novel insights into the differential dynamics of EB3 comets and the microtubule organising centre, demonstrating SReD’s potential for studying complex, time-dependent cellular processes. A key advantage of SReD is its ability to detect and characterise structures without the need for extensive labelled training data or single-molecule localisation input. This feature is particularly useful for exploratory analysis of complex biological systems where the underlying structural patterns may not be fully known *a priori*. The framework’s flexibility in accommodating different reference blocks, from simulated idealised structures to empirically extracted image patches, enhances its utility across diverse experimental scenarios. Another crucial feature is SReD’s robustness to noise and pattern deformations, as demonstrated in our sensitivity analyses. This resilience enables reliable structure detection and quantification even in challenging imaging conditions, expanding the range of biological questions that can be addressed through quantitative image analysis. The algorithm’s multiscale mapping capabilities provide a unique perspective on hierarchical structural organisation, as exemplified by our analysis of nuclear pore complexes at different spatial scales. While SReD offers significant advantages, it is important to acknowledge its limitations. The algorithm’s performance can be influenced by the choice of reference blocks, possibly requiring their optimisation. Additionally, while SReD reduces the need for manual region selection, some level of results curation may still be necessary, particularly in highly complex or heterogeneous samples. Finally, the algorithm’s computational complexity warrants attention. Consider a 2D image with dimensions *n*_1_ × *n*_2_ pixels and a block of size *k* _1_ × *k*_2_ pixels. Each pairwise comparison between the block and an image region requires *O*(*k*_1_ *k*_2_) operations. The total number of such overlapping image regions is (*n*_1_ −*k*_1_ + 1)(*n*_2_ − *k*_2_ + 1). Consequently, the “1-to-all” scheme (block repetition) entails a computational complexity of *O*((*n*_1_ − *k*_1_ + 1)(*n*_2_ −*k*_2_ + 1)*k*_1_*k*_2_). When the image dimensions significantly exceed the block size (*n*_1_ ≫ *k*_1_ and *n*_2_ ≫ *k*_2_), this simplifies to *O*(*n*_1_ *n*_2_ *k*_1_ *k*_2_). In the “all-to-all” scheme (global repetition), the computational complexity scales quadratically with the image size and linearly with the block size, resulting in 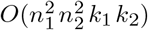. SReD mitigates this computational burden by harnessing GPU acceleration and pre-calculating background regions that do not warrant analysis. Future developments of SReD could focus on further automating the reference block selection process, potentially incorporating machine learning approaches to optimise block parameters based on image characteristics. Integration with other computational tools, such as deep learning-based segmentation algorithms, could also enhance SReD’s capabilities for more comprehensive structural analysis pipelines.

## DATA AVAILABILITY

The data obtained in this study is available at https://doi.org/10.5281/zenodo.13764726 under CC BY 4.0 license. The STORM data containing cells with labelled microtubules is available at (10). The data containing DAPI-stained nuclei is available at (11). The STORM data containing nuclear pores with labelled gp210 is available at (12).

## CODE AVAILABILITY

The SReD algorithm is available at https://github.com/HenriquesLab/SReD. All source code is under an MIT License.

## AUTHOR CONTRIBUTIONS

A.M. and R.H. conceived the study in its initial form; A.M. and R.H. developed the SReD framework with code contributions from B.M.S.; B.M.S., G.J., J.I.M and C.L. provided samples, data, critical feedback, testing and guidance; A.M. performed experiments and analysis; G.J., J.I.M., C.L. and R.H. acquired funding; C.L., R.H. supervised the work; A.M and R.H. wrote the manuscript with input from all authors.

## ACKNOWLEDGEMENTS

A.M. thanks Simão Coelho and Estibaliz Gómez-de-Mariscal for their feedback during the formulation of the algorithm and its demonstration in biological studies. A.M. and R.H. acknowledge the support of the Gulbenkian Foundation (Fundação Calouste Gulbenkian), the European Research Council (ERC) under the European Union’s Horizon 2020 research and innovation programme (grant agreement No. 101001332) (to R.H.) and funding from the European Union through the Horizon Europe program (AI4LIFE project with grant agreement 101057970-AI4LIFE and RT-SuperES project with grant agreement 101099654-RTSuperES to R.H.). Funded by the European Union. Views and opinions expressed are, however, those of the authors only and do not necessarily reflect those of the European Union. Neither the European Union nor the granting authority can be held responsible for them. This work was also supported by a European Molecular Biology Organization (EMBO) installation grant (EMBO-2020-IG-4734 to R.H.), a Chan Zuckerberg Initiative Visual Proteomics Grant (vpi-0000000044 with https://doi.org/10.37921/743590vtudfp to R.H.) and a Chan Zuckerberg Initiative Essential Open Source Software for Science (EOSS6-0000000260). This study was also supported by the Research Council of Finland (338537 to G.J.), the Sigrid Juselius Foundation (to G.J.), the Cancer Society of Finland (Syöpäjärjestöt; to G.J.), the Solutions for Health strategic funding to Åbo Akademi University (to G.J.), the InFLAMES Flagship Programme of the Academy of Finland (decision numbers: 337530, 337531, 357910, and 357911) and the CNRS ATIP program (AO2016 to C.L.). This research was also supported by the National Institutes of Health (NIH) with grants K22AI140963, K61DA058348 and subcontract R01AI50998 (to J.I.M).

## EXTENDED AUTHOR INFORMATION

- A. Mendes: afonsomendes92
- B. Saraiva: Bruno_MSaraiva
- G. Jacquemet: guijacquemet
- J. I. Mamede: Micromede
- C. Leterrier: christlet
- R. Henriques: HenriquesLab

## Methods

### Noise variance stabilisation

The Generalised Anscombe Transform (GAT) is applied to the input image to stabilise noise variance, a crucial step in processing microscopy images. These images typically exhibit a combination of Poisson and Gaussian noise, which can obscure the underlying signal. In fluorescence microscopy, noise variance is signal-dependent, limiting some of the assumptions required for the proper application of correlation metrics. Namely, the assumptions of normality, homoscedasticity (equal variance), and absence of outliers. The GAT employs a nonlinear remapping of pixel values, resulting in an output image with near-Gaussian noise and stabilised variance, whilst preserving local contrast and overall image statistics (9). This variance stabilisation thus improves downstream processing, enabling more accurate analysis of the image’s structural content.

### Relevance mask

We use a relevance mask to filter out areas lacking significant structural information. The rationale is that structural elements present themselves as regional image textures with non-zero variance. Therefore, areas devoid of structure will exhibit minimal texture. Determining the threshold for “minimal” texture is challenging due to the presence of ubiquitous image noise. Rather than choosing an arbitrary value close to zero, we estimated the average noise variance of the input image using a robust estimator (21). This is obtained by sampling the variance of the input image using non-overlapping blocks of the same size as those used for the repetition analysis, and then calculating the average of percentile 0.03. This sampling approach ensures that the noise variance is estimated at the same scale as the analysed structures. The relevance threshold is established by multiplying the estimated average noise variance by an adjustable constant, with the default set at 0. This produces a binary mask outlining areas with sufficient structural content.

### Sampling scheme and mathematical basis

Our algorithm leverages a custom sampling scheme in which a reference block is compared with all possible test blocks in the image. The scheme can be “1-to-all” or “all-to-all”, depending on the application. The first requires a user-provided reference block, while the latter provides unbiased structure detection. The comparisons between blocks consist of calculating correlation metrics. The correlation metrics can be rotation-variant (e.g., Pearson’s correlation coefficient) or invariant (e.g., modified cosine similarity). In both cases, the blocks’ dimensions are predefined by the user to match a specific scale. After defining the blocks’ dimensions and the relevance threshold, the algorithm calculates the noise variance-stabilised input, the relevance mask, and normalises the input image to its range. Then, it calculates the local means and standard deviation maps for the entire universe of blocks that can be extracted from the image within the previously defined constraints. To minimise blocking artefacts, the square blocks are transformed into their inbound elliptical counterparts. Using these statistics, a repetition map is calculated for each “1-vs-all” comparison, where each pixel is assigned a score (named Structural Repetition Score, or SRS), which reflects the similarity between the local neighbourhood centred at that position and the reference block. Finally, the repetition map is normalised to its range. The SRS is given by:

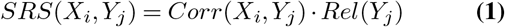

where

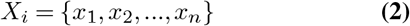

and

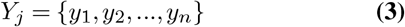

are the reference and test blocks with size *n* (in pixels) centred at pixel positions *i* and *j*, and

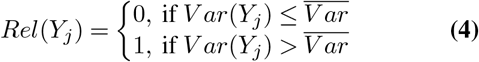

the binary “relevance” label of the test block, where Var is the average noise variance of the input image. To analyse local textures and calculate a single value for each, the reference and test blocks require a defined centre. Therefore, the blocks’ dimensions need to be odd, and as a result, *i* = [(*r*_*h*_, *H* ™ *r*_*h*_] and *j* = [*r*_*w*_, *W* ™ *r*_*w*_], where *r*_*w*_ and *r*_*h*_ are the blocks’ width and height radii, and *W* and *H* are the input image’s width and height. In the global repetition mode, SReD enables unbiased structure analysis by using the entire universe of image blocks as a reference. Each reference block generates a repetition map that is averaged, and the average value is plotted at the coordinates corresponding to the centre of the reference block. The average uses an exponential weight function based on the distance between the standard deviations of the blocks in each comparison, which enhances structural details. Therefore, the global repetition scores represent the relative repetition of a local texture across the image. Mathematically, the global SRS is given by:

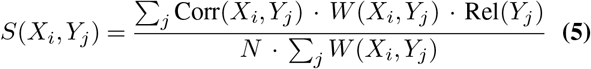

where *N* is the size of the input image (excluding borders), and

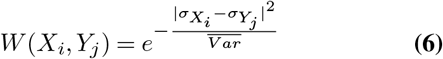

is the exponential weight function, where 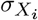 and 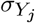 are the standard deviations of the reference and test blocks.

### Multiscale analysis

The scale at which structures are analysed can be adjusted by modulating the ratio between the input image and the block size. For example, larger blocks contain information about higher-order structures compared to smaller blocks. Due to the iterative nature of the algorithm, increasing the block size adds an exponential amount of data points to each comparison, introducing an unwanted load on the computational resources and drastically slowing the calculations. Therefore, modulation of the ratio between the input image and the block size can instead be achieved by adjusting the input image size (i.e., downscaling). A direct consequence of this method is the loss of lower-order information, which should not be problematic when the goal is to increase the sensitivity to higher-order textures.

### Non-linear mapping

Non-linear mapping can enhance the contrast between different SRSs within the repetition maps, facilitating visual interpretation and subsequent analysis. In our experience, we have found that applying a power transformation to the SRSs often yields the most effective enhancement. This transformation involves raising each SRS value to a specific exponent. The choice of exponent plays a crucial role in determining the degree of contrast enhancement. In this study, we explored a range of exponents between 10 and 10,000. Typically, we initiate the analysis with an exponent of 10 and iteratively adjust it based on the visual assessment of the resulting repetition map. For datasets with subtle structural repetitions or low signal-to-noise ratios, higher exponents may be necessary to amplify the differences between SRSs and reveal hidden patterns. Conversely, for datasets with prominent structural repetitions, lower exponents may suffice to achieve adequate contrast enhancement without introducing excessive noise amplification. The optimal exponent ultimately depends on the specific characteristics of the data and the desired level of visual clarity. By carefully selecting the exponent, users can tailor the contrast enhancement to their needs, facilitating the identification and interpretation of repetitive patterns in diverse microscopy images.

### Optimisation of block parameters for ring pattern detection

A collection of 248 testing blocks incorporating various combinations of inter-ring spacing and ring height was generated. This process was automated using a custom ImageJ macro. To create input images, five representative segments from the distal axons within each dataset were randomly selected, comprised of six neurons per treatment. These regions were then rotated to align with the horizontal axis to guarantee consistency across subsequent calculations. Then, SReD was used to generate repetition maps for every test block, and their autocorrelation functions were calculated. The relative amplitude of the autocorrelations’ first harmonic was used to assess how effectively each block captured the underlying periodic pattern. The set of block parameter values that maximised the first harmonics’ relative amplitude was systematically identified. The optimised set of parameter values served as a reliable representation of the periodic pattern within the dataset. The optimisation was performed separately for each dataset analysed in this study.

### Detection of reference and virus-like particles using Global Repetition

An image volume containing 3D simulated reference particles was generated using a custom Python script. The reference particles were added to the input volume by addition. Global Repetition maps were calculated using a block size of 5×5×5 pixels and a relevance constant of 0. Then, the repetition maps were non-linearly mapped using a power transformation with an exponent of 10000. 3D maxima were calculated using the ImageJ “3D Maxima Finder” plugin, with an XY and Z radius of 5 pixels and a minimum threshold of 0.1. The comparison of coordinates between the 3D maxima calculated and the reference particles was performed using a custom Python script.

### Cell culture

Jurkat cells were cultured in RPMI 1640 (Gibco) supplemented with 10% fetal bovine serum (FBS), 2 mM L-glutamine and 50 µg/mL gentamycin. HEK293T and RPE1-EB3-GFP cells were cultured in DMEM supplemented with 10% fetal bovine serum (FBS), 2 mM L-glutamine and 50 µg/mL gentamycin. Cell lines were cultured at 37ºC and 5% CO2.

### DNA plasmids and cell lines

The RPE1-EB3-GFP cell line was kindly provided by Dr. Mónica Bettencourt-Dias. A plasmid expressing HIV-1 Gag with an internal EGFP tag was generated using the NEB HiFi Assembly Kit (New England Biolabs). A lentiviral backbone containing a tetracycline-inducible promoter and a gene encoding rtTA was prepared by digesting the pCW57.1 plasmid (Addgene #41393) with 5 µg/mL restriction enzymes BamHI and NheI (New England Biolabs) according to the manufacturer’s instructions for 1 hour at 37ºC. The digestion product was separated using 1% agarose gel electrophoresis (AGE) and the 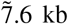 band was purified using the GFX PCR & Gel Band Purification Kit (Sigma-Aldrich) according to the manufacturer’s instructions. Then, three DNA fragments were generated by polymerase chain-reaction (PCR) using Q5 High-Fidelity DNA Polymerase (New England Biolabs). The first fragment (445 bp), encoding the HIV-1 Matrix protein followed by an HIV-1 protease cleavage site (MA-PCS), was generated using Optigag-mNeonGreen-IN (22) as a template and primers *5’-tcagatcgcctggagaattgggccaccatgggtgcgcga-3’ (Fw)* + *5’-ccatacgcgtctggacaatggggtagttttgactgacc-3’ (Rv)*. The second fragment (751 bp), encoding EGFP, was generated using HIV-(i)GFP ΔEnv (18) as a template and primers *5’-ccattgtccagacgcgtatggtgagcaag-3’ (Fw)* + *5’-tagttttgacttctagacttgtacagctcgtc-3’ (Rv)*. The third fragment (1.2 kb), encoding a PCS and the HIV-1 Capsid, Nucleocapsid and p6 proteins (PCS-CA-NC-p6), was generated using Optigag-mNeonGreen-IN (22) as a template and primers *5’-caagtctagaagtcaaaactaccccattgtc-3’ (Fw)* + *5’-aaaggcgcaaccccaaccccgtcattgtgacgaggggtctgaac-3’ (Rv)*. The three fragments were purified using DNA purification columns and their molecular size was confirmed by AGE. The HiFi Assembly reaction was performed using 50 ng of digested vector and equimolar amounts of the three fragments, and incubated at 50ºC for 1 hour. The reaction product was diluted 1:4 in dH20, and 2 µL of the dilution was transformed into chemically competent STABL4 bacteria (Thermo Fisher). The bacteria were plated in LB-agar supplemented with 100 µg/mL ampicillin and incubated overnight at 37ºC. Several colonies were picked and inoculated into liquid LB containing ampicillin at 100 µg/mL. The plasmid DNA from these colonies was extracted using the GenElute Plasmid Miniprep Kit (Sigma-Aldrich), and was confirmed by digestion with restriction enzyme XbaI followed by AGE (2.3 kb and 7.5 kb fragments). A positive colony was then sequenced using Sanger sequencing (Genewiz) and primers *5’-cgtcgccgtccagctcgacca-3’, 5’-ccattgtccagacgcgtatggtgagcaag-3’* and *5’-aaaggcgcaaccccaaccccgtcattgtgacgaggggtctgaac-3’*. This process yielded the lentiviral plasmid TetOn-Optigag-(i)EGFP, where a human codon-optimised *gag* gene contains a PCS-flanked EGFP-encoding gene. Lentivirus packaging TetOn-Optigag-(i)EGFP were produced to transduce Jurkat cells. To do this, HEK293T cells were cultured in 6-well plates until ∼80% of confluence, transfected using 300 µL/well of transfection mixture (DMEM, 3 µg of TetOn-Optigag-(i)EGFP, 1.5 µg of psPAX2 (Addgene #12260), 1.5 µg of CMV-VSV.G (NIAID) and 12 µL of linear polyethyleneimine MW-25,000 (final concentration of 5 µg/µL)(Sigma-Aldrich)) and incubated overnight for 8 hours. Then, the culture medium was replaced with complete DMEM, followed by a 24-hour incubation. The virus-rich supernatant was collected and filtered with 0.22 µm syringe filters. Jurkat cells (2 mL at 1×10^6^ cells/mL) were inoculated with 300 µL of virus-rich supernatant and Polybrene (10 µg/mL), followed by a 3-day incubation. Antibiotic selection of transduced cells was performed by replacing the culture medium with complete RPMI containing puromycin at 2 µg/mL and incubating for 3 days, at which point an “empty virus” control sample had no live cells remaining. The cells were incubated with doxycycline at 1 µg/mL for 24 hours to induce expression and single cells were isolated using Fluorescence-assisted Cell Sorting (FACS). The EGFP-positive population was divided into three subsets according to their relative signal intensity (“Low”, “Medium” and “High”) and single cells were plated into 96-well plates. The cultures were expanded for 15 days and the resulting cell lines were validated using fluorescence microscopy and Western blotting. A clonal line of the “Medium” subset was used for this study.

### Sample preparation and acquisition of microscopy data

#### HILO imaging of HIV-1 virus-like particle assembly in activated Jurkat cells

Activation surfaces were prepared based on the protocol in (Ashdown et al.). To do this, Lab-Tek 8-well chambers (Thermo Fisher) were cleaned with 100% isopropanol for 10 min and followed by three washing steps with dH_2_0. Then, 200 µL of a mouse anti-CD3 antibody diluted in PBS at a final concentration of 1 µg/mL was added to the wells and incubated overnight at 4ºC. The wells were carefully washed twice with PBS to remove unbound antibodies. Jurkat cells expressing TetOn-Optigag-(i)EGFP were incubated with 1 µM of doxycycline (Sigma Aldrich) for 24 hours. Then, 50000 cells were added to each well and allowed to adhere and stabilise for 1 hour. Imaging was done in a Nanoimager (ONI) using the 488 nm laser at 10% and channel 0 (two-band dichroic: 498-551 nm and 576-620 nm). The HILO angle was optimised manually and images were acquired at 100 ms exposure. The anti-CD3 antibody was produced at the Flow Cytometry & Antibodies Unit of Instituto Gulbenkian de Ciência, Oeiras, Portugal.

#### 3D imaging of HIV-1 virus-like particle assembly in activated Jurkat cells

Jurkat cells expressing TetOn-Optigag-(i)EGFP were centrifuged at 200 xg for 5 minutes and resuspended in complete RPMI containing 0.5 µM of doxycycline to induce Gag expression. Glass coverslips (1.5 mm thick, round, 18 mm diameter) were washed with isopropanol for 10 minutes followed by three washes with dH_2_0. The coverslips were coated with Poly-L-Lysine (PLL, Sigma Aldrich) at 0.1% and incubated for 15 minutes at room temperature, followed by three washing steps with dH_2_0. The PLL-coated coverslips were dried, mounted in an Attofluor chamber (Thermo Fisher) and fixed on the microscope’s stage. The microscope’s enclosure (Okolabs) was heated at 37^°^C and a manual gas mixer (Okolabs) was used to supply 5% CO_2_. The cells were seeded in the pre-treated coverslips and allowed to settle in the microscope enclosure for 30 minutes. Imaging was performed on an inverted microscope ECLIPSE Ti2-E (Nikon Instruments) equipped with a Fusion BT (Hamamatsu Photonics K.K., C14440-20UP) and a Plan Apo *λ* 100x (NA 1.45) Oil objective. The sample was illuminated with LED light at 515 nm (CoolLED pe800) and acquisition was done at 75 ms exposure with an active Nikon Perfect Focus system and the NIS-Elements AR 5.30.05 software (Nikon Instruments). Volumes were captured by acquiring frames at different depths (z-step size: 0.5 µm). Image deconvolution was performed using a custom Python script based on the Richardson-Lucy method (23, 24) as described in (25, 26).

#### Imaging of EB3-GFP comets in RPE1 cells

RPE1-EB3-GFP cells (50000 per well) were seeded into Lab-Tek 8-well glass chambers (Thermo Fisher) and allowed to adhere for 24 hours. Imaging was performed in a Nanoimager (ONI) using the 488 nm laser at 10% and channel 0 (two-band dichroic: 498-551 nm and 576-620 nm). Images were acquired at 75 ms exposure for 2 minutes.

### Assessment of microtubule dynamics using SReD

Subsets of the original time lapse were created by keeping images belonging to the time frames of interest. Global repetition maps were generated from the temporal subsets using an XY block size of 7×7 pixels, a Z block size equal to the number of images in each subset, and a relevance constant of 0. The repetition maps were non-linearly mapped using a power transformation with an exponent of 1000. NRMSE maps were calculated using the “scikit-image” library (v0.22.0).

## Note S1: Theoretical foundation and core functionality of SReD

The Structural Repetition Detector (SReD) is an unsupervised computational framework designed to identify repetitive biological structures in microscopy images by exploiting local texture repetition. SReD operates by comparing local image regions (blocks) to detect recurring patterns without prior knowledge or constraints on the imaging modality. The algorithm’s workflow includes the following preprocessing steps:

1. **Application of the Generalized Anscombe Transform (GAT):** This step stabilises noise variance, addressing the complex noise characteristics typical in microscopy images. The GAT employs a nonlinear remapping of pixel values to produce an output image with near-Gaussian noise and stabilised variance, preserving local contrast and overall image statistics (1). The pixel values of the transformed image are given by:

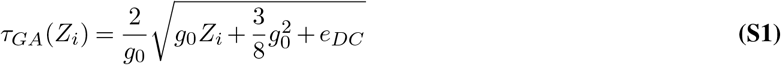

where *g*_0_ is the gain of the electronic system, *e*_*DC*_ = *σ*^2^− *g*_0_*m*, and *m* and *σ*^2^are the mean and variance of the noise. The goal is to find parameter values such that the transformed image has variance as close as possible to 1. The initial parameter values can be user-provided or calculated automatically.
2. **Generation of a Relevance Mask:** This mask excludes regions lacking substantive structural information, based on local texture prominence quantified by variance or standard deviation. The relevance threshold is defined by multiplying the estimated average noise variance (2) by an adjustable constant, with the default set at 0.

SReD’s primary functionality revolves around two main analysis modes:

1. **Block Repetition Analysis:** The input image is probed for repetitions of a single reference block, which can be either simulated or empirically extracted from the data. The output is a repetition map reflecting the likelihood of the reference pattern occurring at each location. The similarity score is computed using a correlation metric, which can be sensitive or insensitive to rotation.
2. **Global Repetition Analysis:** Every block in the input image is used as a reference, generating multiple repetition maps. These maps are integrated using an exponentially weighted average based on block similarity, producing a final map that reflects the relative frequency of each structural pattern across the entire image.

The algorithm’s output is a Structural Repetition Score (SRS) for each pixel, quantifying the degree of local structural repetition. This score can be interpreted as the likelihood of a specific pattern occurring at that location. SReD also offers multiscale analysis capabilities by adjusting the ratio between the input image and block sizes, enabling the detection of structural patterns at various scales. This approach is particularly valuable for exploratory analysis of complex biological systems where the underlying structural patterns may not be fully known a priori.

## Note S2: Validation and application examples of SReD across diverse biological contexts

### Detection of microtubules using simulated reference blocks

We evaluated SReD’s ability to detect structures using simulated reference blocks. A series of blocks comprising lines and line crossings at different orientations were generated and used to detect microtubule structures in a STORM image reconstruction of a HeLa cell with labelled microtubules (3) (**Fig. S2a**). Using the Pearson’s correlation coefficient metric for orientation sensitivity, SReD produced repetition maps highlighting regions matching the simulated blocks (**Fig. S2b**). The specificity was assessed by:

1. Detecting repetitions of a vertical line block in images with vertical lines at varying distances
2. Detecting repetitions of an orthogonal line crossing block in images with crossings at different angles

In both cases, the average SRS decreased markedly when the specific structures were absent, demonstrating high specificity (**Fig. S2c,d**).

### Detection of nuclear envelopes using empirical reference blocks

To demonstrate structure detection using empirical reference blocks, we extracted blocks containing nuclear envelope regions at various orientations from a DAPI-stained image of HeLa cell nuclei (4) (**Fig. S3a**). Applying SReD with the Pearson correlation metric generated repetition maps highlighting regions matching the reference blocks, effectively mapping the nuclear envelopes and their local orientations (**Fig. S3b,c**). This approach also enables the characterisation of morphological variations. We calculated the relative area percentage of each repetition map in the combined output to quantitatively describe nuclear shape (**Fig. S3d**). These measurements distinguished different morphological states potentially related to cell division or stress.

### Detection of HIV-1 virus-like particles using Global Repetition

SReD’s global repetition mode enables unbiased structure detection by quantifying the relative repetition of all structures in an image. We analysed an image of a Jurkat cell expressing an HIV-1 Gag-EGFP construct that induces virus-like particle (VLP) assembly (**Fig. S4a,b**). The global repetition map, calculated using modified cosine similarity, showed:

1. The most repetitive structure was the cell background signal (diffusing Gag protein)
2. Lower SRS structures corresponded to active VLP assembly sites
3. Some structures not discernible in the input image were detected, likely early VLP assembly stages

We identified VLP locations by calculating local maxima. The repetition map provided more detections than the input image when using the same prominence threshold (**Fig. S4c**).

### Multiscale detection of Nuclear Pore Complex structures

SReD’s multiscale analysis capability was demonstrated by examining nuclear pore complexes (NPCs) in STORM reconstructions of labelled gp210 proteins. Global repetition maps were calculated at different scales by modifying block-to-image size ratios (**Fig. S5a,b**):

1. Original image (400×400 pixels), 5×5 pixel blocks: detected single nucleoporins
2. Original image, 15×15 pixel blocks: detected nucleoporin clusters
3. Downscaled image (200×200 pixels), 25×25 pixel blocks: detected entire NPC units

This demonstrates SReD’s ability to detect structures at multiple scales without requiring structural priors.

## Note S3: Automated analysis pipeline for spectrin ring detection in neuronal axons

### Automatic estimation of axon orientations

A robust analysis of ring periodicity using autocorrelation functions requires the axon segments to be oriented with their long axis parallel to the horizontal axis. We designed a streamlined approach that uses SReD’s block repetition to estimate the orientation of the axons along their span (**Fig. S6a**). This information can be used for downstream applications, such as estimating the distribution of angles in each sample (**Fig. S6b,c**) or automatic region extraction and rotation. Our method is fully implemented in ImageJ macros and works as follows:

1. **Calculate the axons’ “skeletons”**. “Skeletonize” function produces a binary mask of the axons’ skeletons by shrinking their to a 1-pixel wide line along their centers of mass. To improve the results of this function, the images are preprocessed by applying a Gaussian blur (15 pixel radius) to retrieve only the high-order structures and discard the single-molecule low-order information. Then, thresholding is performed using the Otsu method (5) to remove unwanted objects. The “Skeletonize” function is applied to the thresholded images. A Gaussian blur (2 pixel radius) is applied to the skeletons to avoid diagonal discontinuities derived from the 1 pixel-wide lines that form the skeletons. Finally, a range normalisation step is performed to bring all the skeletons to the same intensity interval.
2. **Generate synthetic blocks comprising lines at different orientations**. This is done by designing a block containing a vertical 1 pixel-wide line in a 90×90 black canvas. Then, copies of this block are created and rotated in 10º increments, until all possible orientations are recapitulated at that angular resolution. A Gaussian blur (2 pixel radius) is applied to the blocks to match the appearance of the blurred axon skeletons. The synthetic blocks are then cropped into 45×45 pixel blocks to avoid border artefacts, and normalised to their intensity range.
3. **Calculate block repetition maps using SReD**. SReD’s block repetition is used to generate repetition maps using the synthetic blocks and axon skeletons generated previously. Each round of block repetition produces a repetition map where regions containing skeletons at the specific orientations display higher SRS. The repetitions maps are normalised to their intensity range, multiplied by the Gaussian blurred masks to remove unwanted detections, and normalised once more.
4. **Generate the angle maps**. Each coordinate of the 1 pixel-wide skeletons is labelled with the angle corresponding to the highest SRS in the repetition maps.

### Reference block optimisation for ring pattern detection

The detection of specific structures requires the use of a reference. In SReD, the reference is provided as an image block containing a representation of the structure of interest. To minimise the bias of downstream analyses towards the reference block, we devised a method to optimise the reference block based on the data characteristics (**Fig. S7a**). The method is implemented as a combination of ImageJ macros and Python code. It works as follows:

1. **Conceptualisation of the reference block**. The reference block used for spectrin ring detection should recapitulate a periodic ring pattern while using the minimum amount of information to minimise bias towards that pattern’s characteristics. We concluded that the simplest pattern would comprise a black canvas with 3 vertical lines resembling a side-view of a 2D projection of rings. While using 2 lines instead of 3 seems more parsimonious, doing so would centre the pattern at the inter-ring spacing and not the ring itself. The 3-ring pattern can be defined by two parameters: (i) inter-ring spacing and (ii) ring height. The first parameter is the main target of our study, while the latter is less important and is mostly a function of axon girth.
2. **Optimisation of the ring pattern’s parameters**. To minimise the bias of our study towards the reference pattern, a parameter sweep was performed to optimise the pattern’s characteristics according to the input data. To do this, we generated a collection of 248 blocks incorporating various combinations of inter-ring spacing and ring height. Representative segments of distal axons (6 for each group) were extracted from the data. From these axon segments, a total of 30 test regions were extracted and rotated to align with the horizontal axis. This rotational adjustment guaranteed consistency when applying the same set of reference blocks across the datasets, eliminating any potential variations stemming from block rotation and interpolation. SReD was used to calculate block repetition maps for every reference block and the autocorrelation functions of the repetition maps was calculated. The relative amplitude of the autocorrelations’ first harmonic was used to assess how effectively each block captured the underlying periodic pattern. We systematically identified the set of block parameter values that maximised the first harmonic’s relative amplitude (**Fig. S7b-d**). This optimised set of parameter values serves as a reliable representation of the periodic pattern within the dataset. The optimisation was performed separately for each dataset analysed in this study.

### Detection of ring patterns in large fields-of-view

Using the optimised ring pattern and the angle distributions previously generated, SReD can be used to map ring patterns at all orientations across large fields-of-view.

1. **Preparing the input data for analysis** An input image comprising a large-field-of-view of axons featuring labelled spectrin rings is copied and rotated for each angle defined previously (0º to 180º in 10º steps). To avoid cropping at the borders, the image is zero-padded before this step. A range normalisation is also applied.
2. **Detecting spectrin rings at all orientations**. Block repetition maps are calculated for each rotated input. Doing this ensures that each axonal region in the input is probed at least once while oriented parallel to the horizontal axis. Then, the repetition maps are rotated back to their original orientation.
3. **Generation of angular weight maps** From the repetition maps calculated previously, we are interested in retrieving only the information that is relevant for a specific angle. For example, in the repetition map calculated from the input image rotated by -10º, we want to keep the segments whose orientation relative to the horizontal axis was estimated to be 10º. To do this, we use a weight multiplication approach. An angular weight map is generated for each orientation. The weight maps are generated using the angle repetition maps calculated previously. The maps are slightly transformed to accommodate the extent of the axons in the input data by applying a power function (exponent of 8) and a Gaussian blur (radius 2 pixels), followed by a range normalisation step.
4. **Application of the angular weight maps and generation of the final output** The angular weight maps are multiplied by the ring repetition maps that were calculated using inputs rotated to their corresponding angle. For example, the ring repetition map calculated from the input image rotated 10º is multiplied by the angular weight map comprising regions that are oriented at 10º from the horizontal axis. This step retains only the ring pattern information that is relevant at each orientation. After a range normalisation step, the final image reconstruction is generated by averaging the weighted repetition maps (**Fig. 2b**).

### Automatic extraction and quantitative analysis of ring patterns in axon segments

Previous studies focusing on characterising the periodicity of spectrin rings in axons were limited by the need to manually select regions for analysis. This method is labour-intensive and the results can vary between experts. Semi-automatic methods also exist, where algorithms such as NeuronJ (6) produce tracings of the neurons and provide a framework for quantitative analysis. However, these algorithms do not take into account the underlying low-order structures, resulting in axon segments that may or may not contain periodic ring patterns, which impacts their quantitative descriptions. Our approach leverages SReD’s multiscale capabilities to ensure that only regions containing patterns are analysed:

1. **Detection of high-order ring patterns**. Ensuring that the axon segments analysed contain ring patterns is essential to avoid corrupting the results with unwanted structural patterns. This entails mapping regions of the input data where high-order patterns exist. This is often achieved by convolving data with Gaussian kernels. However, this method inherently discards the low-order information and its results are mostly a function of local signal intensity. We approach this problem by leveraging SReD’s multiscale analysis capabilities. Using the optimised ring reference block, a new reference block comprising an extension of the original is generated with 9 rings instead of 3 (**Fig. S8a**). SReD is used to calculate block repetition maps, in which regions where the high-order patterns are more likely to exist will have higher SRS (**Fig. S8b**). A quantitative comparison between our approach and the more common approach using convolutions with Gaussian kernels demonstrates that SReD produce a more robust analysis by retaining the low-order information of ring units while providing mapping the presence of high-order patterns (**Fig. S8c,d**). The high-order repetition map is thresholded using the Otsu method (5) to generate a binary mask where regions containing high-order patterns are segmented. These regions can be used for downstream analysis using the optimised low-order reference block, ensuring that the data analysed contains the structures of interest.
2. **Removing axon crossings and bifurcations**. This step is essential to the study of axon ring periodicity because (i) regions containing axon crossings may contain multiple periodic patterns at different phases and orientations, and (ii) the arrangement of spectrin in regions where axons bifurcate is not well-characterised and is known to not display the periodic patterns observed in more distal regions (7). These regions are filtered out from the analysis using a custom ImageJ macro that detects coordinates where a pixel of the skeleton contacts with more than 2 pixels. This creates discontinuities in the skeletons exactly where the crossings/bifurcations are detected. Then, the extremities of the isolated segments are iteratively pruned, until each extremity is reduced by an amount of pixels equal to the radius of the regions chosen for downstream analysis (75 pixels). This completely eliminates unwanted regions.
3. **Extraction of regions containing periodic patterns from the input data**. The segmented axon skeletons generated in the previous step are used as a basis to extract structurally relevant regions (**Fig. S9a**). The characteristics of the segments already provide a basis to study the stability of spectrin rings, enabling comparisons between control axons and axons treated with swinholide A. We demonstrate this by calculating their length distribution. No significant differences were found between the length distributions, indicating that swinholide A treatment did not impact the high-order arrangement of spectrin rings (**Fig. S9b**). To automatically extract regions for the analysis of low-order patterns, the axon segments are divided into non-overlapping 150×150 pixel blocks centred at their skeletons. The coordinates of each block are used to crop the corresponding regions from the large field-of-view ring repetition maps generated previously. Then, each block is rotated to become parallel to the horizontal axis using the angle estimations calculated previously. This process yields a library of regions containing ring patterns and their ring repetition maps (**Fig. S9c**).
4. **Quantitative analysis of local ring patterns**. The rings patterns detected and extracted using SReD can be used for downstream analysis. This enables the characterisation of the patterns’ characteristics. We calculated the autocorrelation functions of each region and, from the first harmonics’ position, determined that the average inter-ring spacing in our datasets was approximately 180 nm. We found no significant differences between the control and the swinholide Atreated samples (**Fig. S9d**). The first harmonics’ amplitude can be interpreted as the strength or prominence of the periodic pattern. Here, we found a 12% reduction in prominence in the samples treated with swinholide A (**Fig. S9e**). This reduction was smaller compared to previous studies (7). However, when we analysed the fraction of regions containing periodic patterns, we found a 39% reduction in the samples treated with swinholide A that was not reported previously. These results demonstrate that our approach enabled discerning the effects of swinholide A in pattern frequency from pattern prominence.
5. **Evaluation of SReD’s performance in noisy data**. The image reconstructions used in the previous sections were generated from localisation data obtained using Single-Molecule Localisation Microscopy (SMLM). Due to the synthetic nature of these reconstructions, the images produced are devoid of noise originated from the electronic acquisition systems. However, the generalisability of our method to other microscopy modalities requires sensitivity to structures in noisy data. We evaluate SReD’s performance in noisy data using a test image of an axon segment, to which Gaussian noise with incrementally higher standard deviation is added. SReD is then used to detect repetitions of the optimised ring block and its performance is evaluated by comparing the block repetition maps with the “noise-free” control sample (**Fig. S10a**). The repetition maps calculated showed that SReD was able to detect ring patterns in a wide range of signal-to-noise ratios (SNRs). Notably, it enabled detecting ring patterns in images with very low SNR, where structures are usually not discernible (**Fig. S10b**). Despite the reduction in confidence, evidenced by the lower SRSs, the detections remained specific, as shown by their colocalisation with the reference structures (**Fig. S10b**, bottom row). We quantified the performance of the algorithm by calculating correlation metrics between the noisy inputs and the control (**Fig. S10c**). We used two metrics commonly used to assess image quality and fidelity (the Structural Similarity Index Measure (SSIM) and the Root Means Squared Error (RMSE)). This analysis showed that the repetition maps were a more robust representation of the control sample. We also analysed the results using autocorrelation functions. Here, the autocorrelation harmonics degraded quickly in the noisy input images while remaining well-defined in the repetition maps (**Fig. S10d**). The characteristics of the autocorrelations’ harmonics revealed that the expected inter-ring spacing of approximately 190 nm was discernible in all repetition maps, while in the input data it was only detected in the control sample. The prominence of the periodic patterns attributes high confidence to these conclusions (**Fig. S10e**). Together, our results show that SReD’s repetition maps were a superior platform for the quantitative analysis of periodic patterns and structure detection in noisy data when compared to the direct analysis of input data.
6. **Evaluation of SReD’s specificity**.When detecting repetitions of a reference structure, it is important to evaluate their specificity, since a common caveat of this approach is the introduction of bias towards the reference. This could result in false-positive detections. We evaluated SReD’s specificity by analysing an image of axon segment whose width was incrementally stretched, disrupting the periodic ring pattern’s characteristics (**Fig. S11a**). SReD was used to calculate repetition maps using the optimised reference block (**Fig. S11b**). The average SRS across the repetition maps decreased abruptly upon stretching the input image, indicating a high specificity for the reference structure (**Fig. S11c**). Importantly, this decrease in the average SRS was followed by a slight increase that then faded, suggesting the detection of a secondary pattern when the input image was stretched to twice the original width. This observation was investigated by analysing the corresponding repetition maps. We determined that the secondary pattern corresponded to the detection of the second harmonic of the ring periodic pattern. This was explained by the alignment of the reference block’s centre line with a single spectrin ring or the two outer lines with two spectrin rings (**Fig. S11d,e**). The confidence of this detection was highest at a stretch factor of 2 because the patterns period is exactly twice the original at this level. Furthermore, autocorrelation functions showed that, in these conditions, the peak of the second harmonic colocalised with the input’s intrinsic pattern (**Fig. S11f**). Analysis of the first harmonic of the autocorrelations revealed that the period of the pattern in the input image increased with the stretch factor, whereas in the repetition maps it remained at approximately 180 nm but with incrementally lower prominence, indicating the high specificity of the algorithm. The second harmonic remained at twice the period of the first harmonic while the reference pattern was present, and reflected the input’s intrinsic period thereafter (**Fig. S11g,h**). Together, these results suggest that SReD is highly specific for the reference structures, with false-positive detections still reflecting the structural arrangements of the input data and being discernible from true-positives by the magnitude of the SRS.

## Note S4: Evaluation of SReD’s performance in images corrupted with non-specific structures

### Generating images containing non-specific structures

Due to non-specific labelling and autofluorescence, microscopy data often contains unwanted structures, corrupting its integrity. This presents a challenge for structural analysis. Unlike camera noise, which can be effectively addressed through established denoising algorithms leveraging its predictable characteristics, non-specific structures add complexity by introducing additional structures into the data. This can obscure specific structures or alter their appearance, complicating accurate analysis. While an expert might visually distinguish specific from non-specific structures in certain cases, their presence still impacts the analysis. Given that SReD’s global repetition mode is designed to detect all structures in an image, it was crucial to evaluate how non-specific structures affect the analysis of specific structures in a controlled sample. To achieve this, non-specific structures were introduced into images of a Jurkat cell producing viruslike particles (VLPs), along with simulated free particles exhibiting similar characteristics (**Fig. S12a**). Perlin noise at varying frequencies was added to simulate non-specific structures with different levels of complexity using the Python (3.9.4) “Perlin-Noise” library (v1.13). Different noise frequencies were achieved by modifying the “scale” parameter, which corresponds to the inverse of the frequency. Higher frequencies generate structures with increased complexity.

### Analysis of SReD’s robustness against non-specific structures

Global repetition maps were calculated for each sample (**Fig. S12b**). The coordinates of small round objects were identified by calculating 3D local maxima. The number of detections in the global repetition maps was higher than in the input images for all samples (**Fig. S12c**). Specifically, the input images had detection counts of 251, 235, and 223 at noise frequencies 0, 0.02 and 0.04, while the global repetition maps showed detection counts of 505, 281, and 462 (**Fig. S12d**). Furthermore, the percentage of ground-truth detections was substantially higher in the global repetition maps compared to the input images. In the input images, the ground-truth detection percentages were 32%, 30%, and 24%, whereas in the global repetition maps, these percentages increased to 96%, 63%, and 79%, respectively (**Fig. S12e**). These results underscore the significant impact of non-specific structures on structural analysis and highlight the enhanced detection capabilities of SReD’s global repetition mode. By substantially increasing the detection of ground-truth structures, even in the presence of complex non-specific noise, this method demonstrates its robustness and effectiveness in accurately identifying specific structures within corrupted microscopy data.

## Note S5: Assessment of microtubule dynamics using SReD

### Calculation of spatiotemporal global repetition maps

The versatility of SReD’s global repetition mode is demonstrated by extending its 3D capabilities to enable spatiotemporal analysis. A 2D time-lapse was analysed by defining time as the third dimension, resulting in the analysis of texture repetitions within a given time interval. Several time intervals were defined, and subsets of the input data were created by only keeping images within those intervals. Then, the depth of SReD’s block size was defined as having the same size as the number of images in each subset. Since SReD’s sampling scheme dictates that blocks are not allowed to analyse regions beyond the data’s dimensions, this results in a single repetition map for each subset, which highlights the degree of repetition of local textures across the specified time intervals (**Fig. S13a**). Here, the global SRSs can be interpreted as the spatiotemporal stability of structures at each coordinate, with higher SRSs representing more stable structures and *vice versa*.

### Analysis of microtubule dynamics

Assessment of microtubule dynamics was conducted using spatiotemporal global repetition maps. These maps were generated by comparing the state of structures at each time interval with their initial state, employing NRMSE metrics (**Fig. S13a**). Higher NRMSE values indicated less stability of structures over time, and *vice versa*. NRMSE maps derived from SReD’s repetition maps were compared to maps derived from (i) final images of each time interval and (ii) temporal projections. Visual analysis revealed that NRMSE maps from input images highlighted displacements between successive time points. Some regions with high error values in these maps corresponded to local noise patterns because the block size required to capture structures was unable to capture the average noise distribution. NRMSE maps from temporal projections, due to their additive nature, were substantially corrupted by local noise differences. Consequently, these maps poorly distinguished specific structural stability from noise. In contrast, global repetition maps remained highly robust against local noise variations and effectively emphasised the recurrence of structures over time. The global repetition maps combined the evaluation of structure displacement over time with measurements of structure repetition, providing a superior platform for analysis. In these maps, NRMSE values highlighted regions with low NRMSE, indicating that the underlying structures (MTOCs), were significantly more stable compared to EB3 comets. This superior capability of global repetition maps to detect differences between time intervals is attributed to their resistance to noise corruption and their accurate representation of spatiotemporal information. While the average NRMSE values from input images and temporal projections remained stable over time, SReD effectively captured the dynamic nature of microtubules, showcasing its efficacy in tracking and analysing microtubule dynamics (**Fig. S13b**).

---

**Fig. S1.**
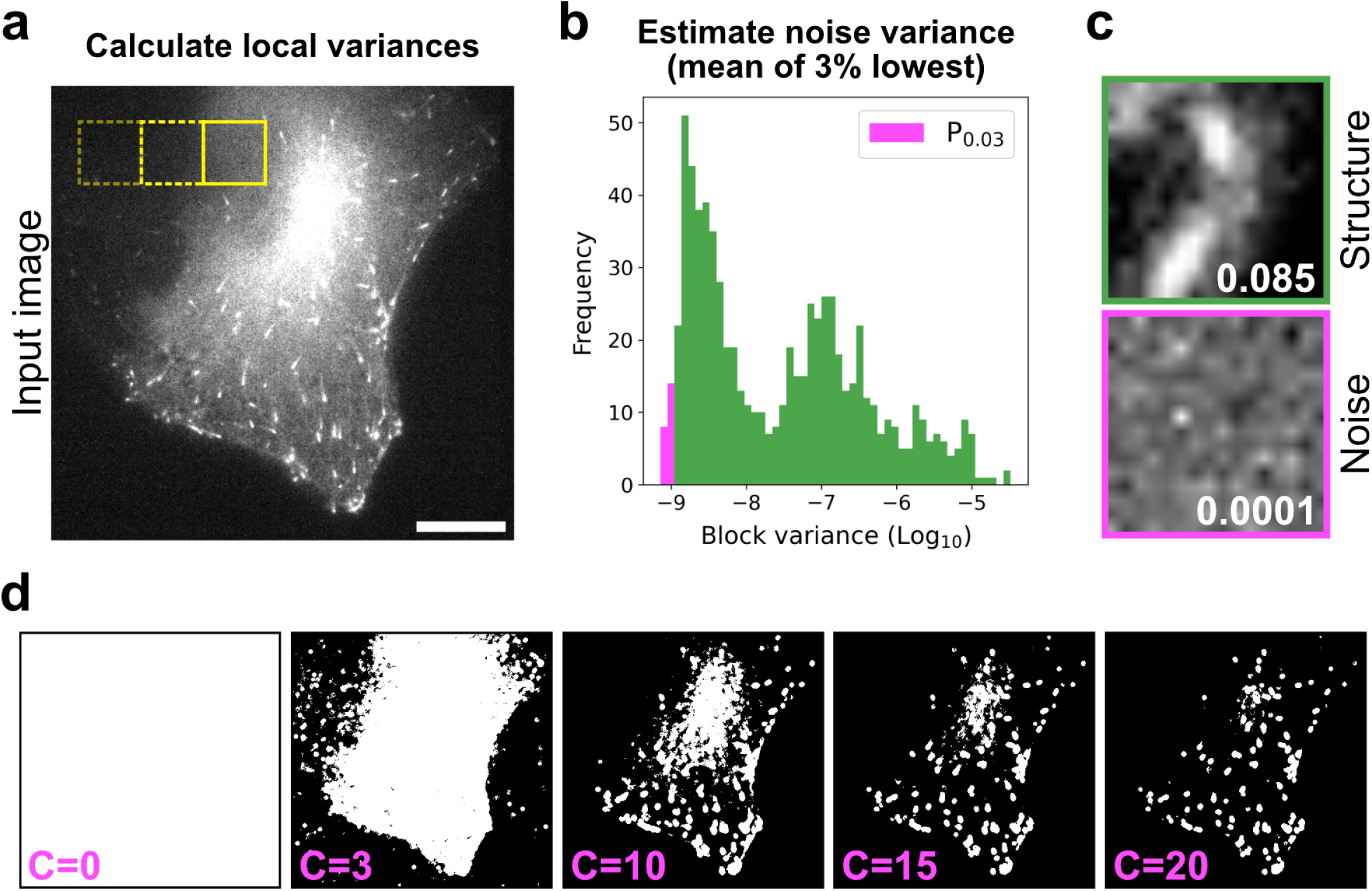
Generation of the relevance mask. **a**, Representative input image with local variance calculated using non-overlapping blocks, illustrating spatial distribution of image texture. Scale bar: 10 µm. **b**, Histogram of calculated block variances, highlighting the lower tail (magenta, percentile 0.03) and upper tail (green) of the distribution. **c**, Example blocks extracted from the distribution tails: upper tail (green border) showing regions with significant structural information, and lower tail (magenta border) representing predominantly noise. The variance of each block is shown in white text. **d**, Series of relevance masks generated using different filter constants (C). Each mask is created by multiplying C with the previously calculated average noise variance to determine the final threshold for structural relevance. This demonstrates how adjusting C impacts the discrimination between structurally relevant and irrelevant image regions.

**Fig. S2.**
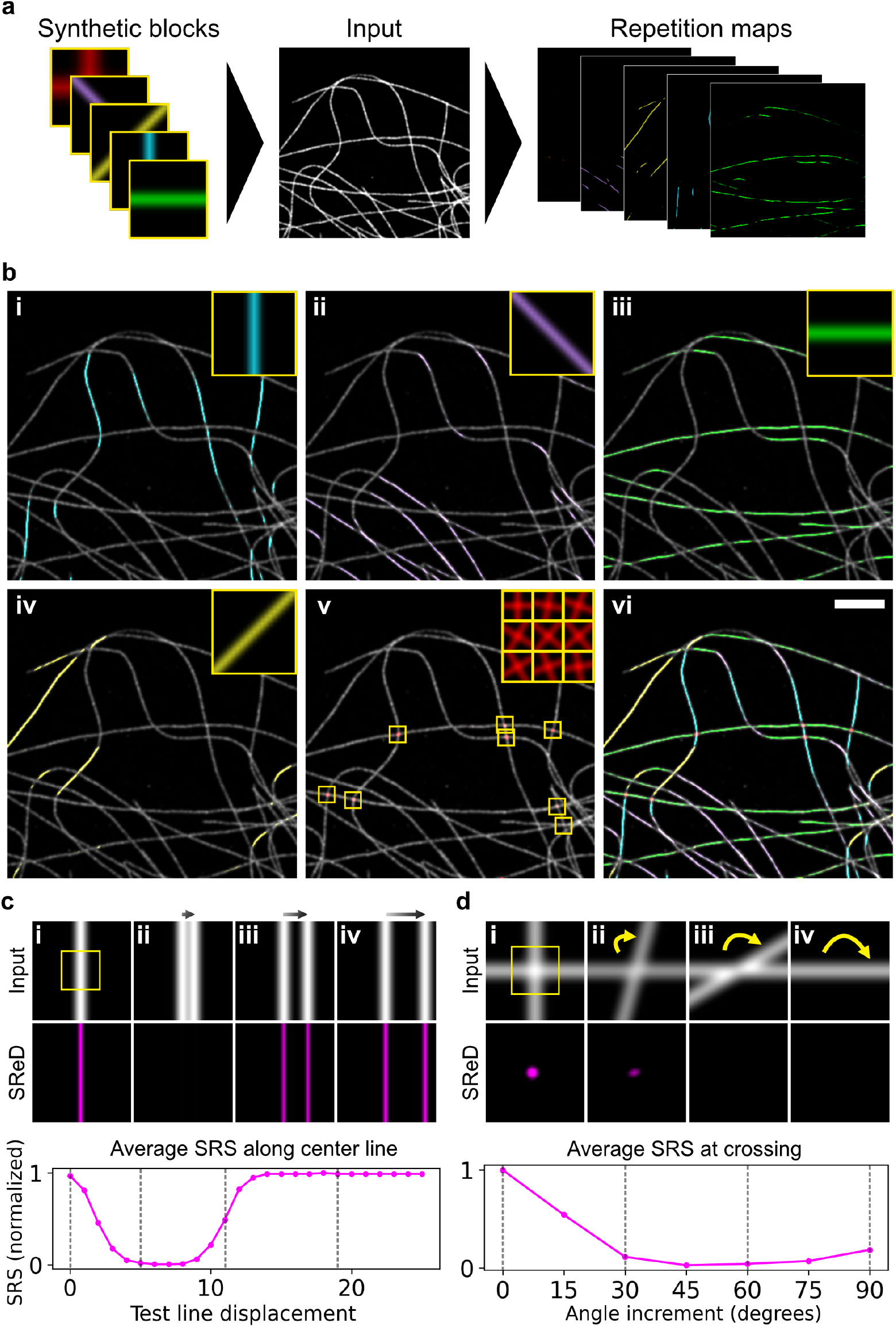
Structure detection using simulated blocks. **a**, Workflow diagram illustrating the use of simulated reference blocks for structure detection. Simulated blocks containing lines and line crossings at various orientations were generated and used by SReD to analyze a STORM image of microtubules in a HeLa cell. SReD produces a repetition map for each reference block, highlighting regions of structural similarity. **b**, Composite images showing the original STORM reconstruction (grey) overlaid with repetition maps (colour) for each simulated reference block. Scale bar: 2 µm. **c**, Analysis of SReD’s specificity for linear structures at varying proximities. (i-iv) Repetition maps generated using a vertical line reference block (yellow box in i) on test images with vertical lines at increasing distances. Graph shows the average Structural Repetition Score (SRS) versus line displacement, with dashed lines corresponding to examples i-iv. **d**, Evaluation of SReD’s specificity for complex structures. (i-iv) Repetition maps generated using an orthogonal line crossing reference block (yellow box in i) on test images with line crossings at increasing angles. Graph displays the average SRS versus angle increment, with dashed lines corresponding to examples i-iv.

**Fig. S3.**
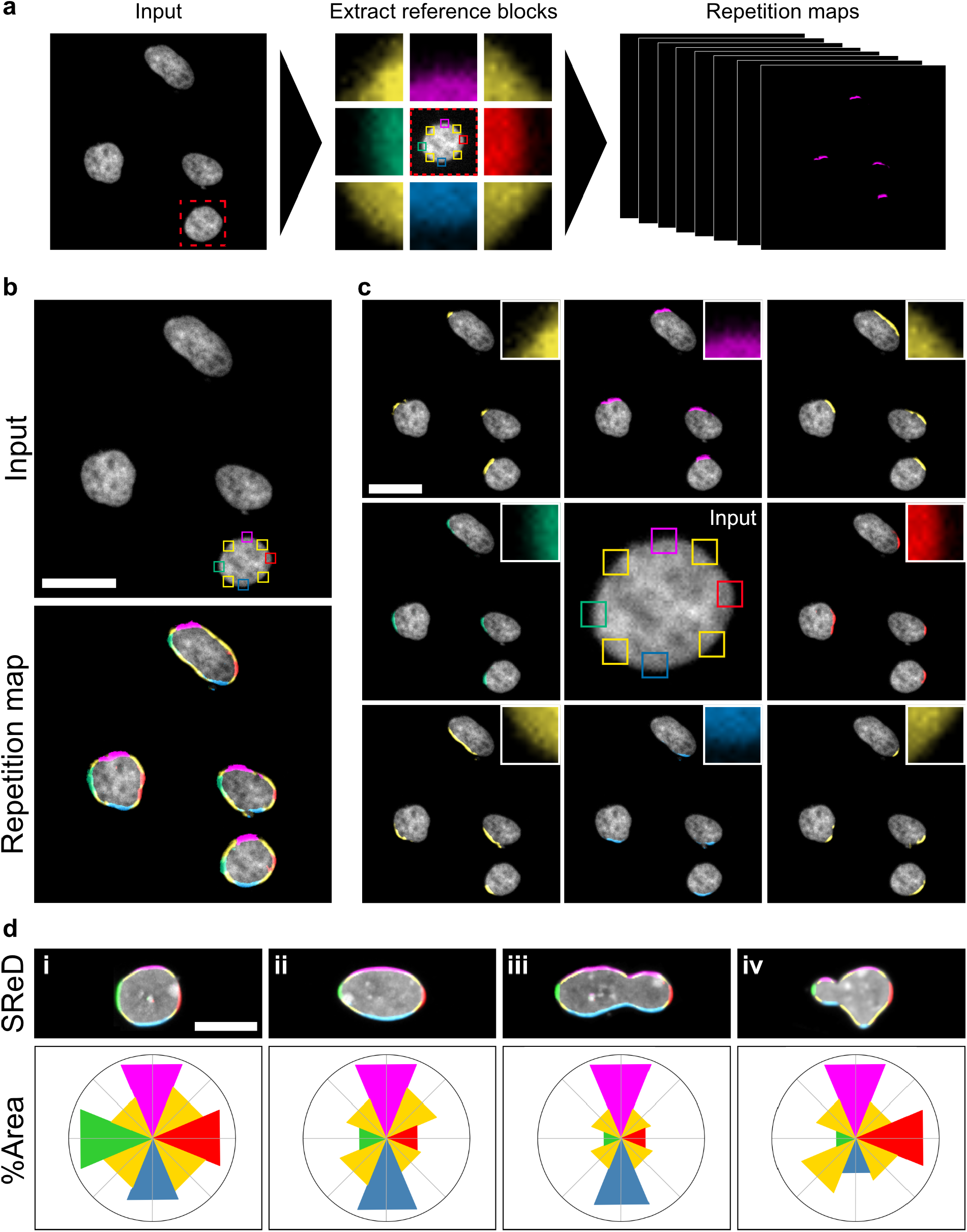
Structure detection using empirical blocks. **a**, Workflow diagram illustrating the use of empirically extracted reference blocks for structure detection. Reference blocks containing nuclear envelope regions at various orientations were extracted from a DAPI-stained image of HeLa cell nuclei. SReD generates a repetition map for each reference block, highlighting regions of structural similarity. **b**, Top panel: Input image of DAPI-stained HeLa cell nuclei with locations of extracted empirical blocks highlighted. Bottom panel: Composite image showing the overlay of colour-coded repetition maps, each corresponding to a different empirical reference block. Scale bar: 30 µm. **c**, Expanded view of individual repetition maps for each empirical reference block, demonstrating SReD’s ability to detect nuclear envelope structures at different orientations. Scale bar: 30 µm. **d**, Top panels: Composite images showing the overlay of colour-coded repetition maps of nuclei i-iv. Scale bar: 30 µm. Bottom panels: Polar plots depicting the relative area percentages of each repetition map in the nuclei composites.

**Fig. S4.**
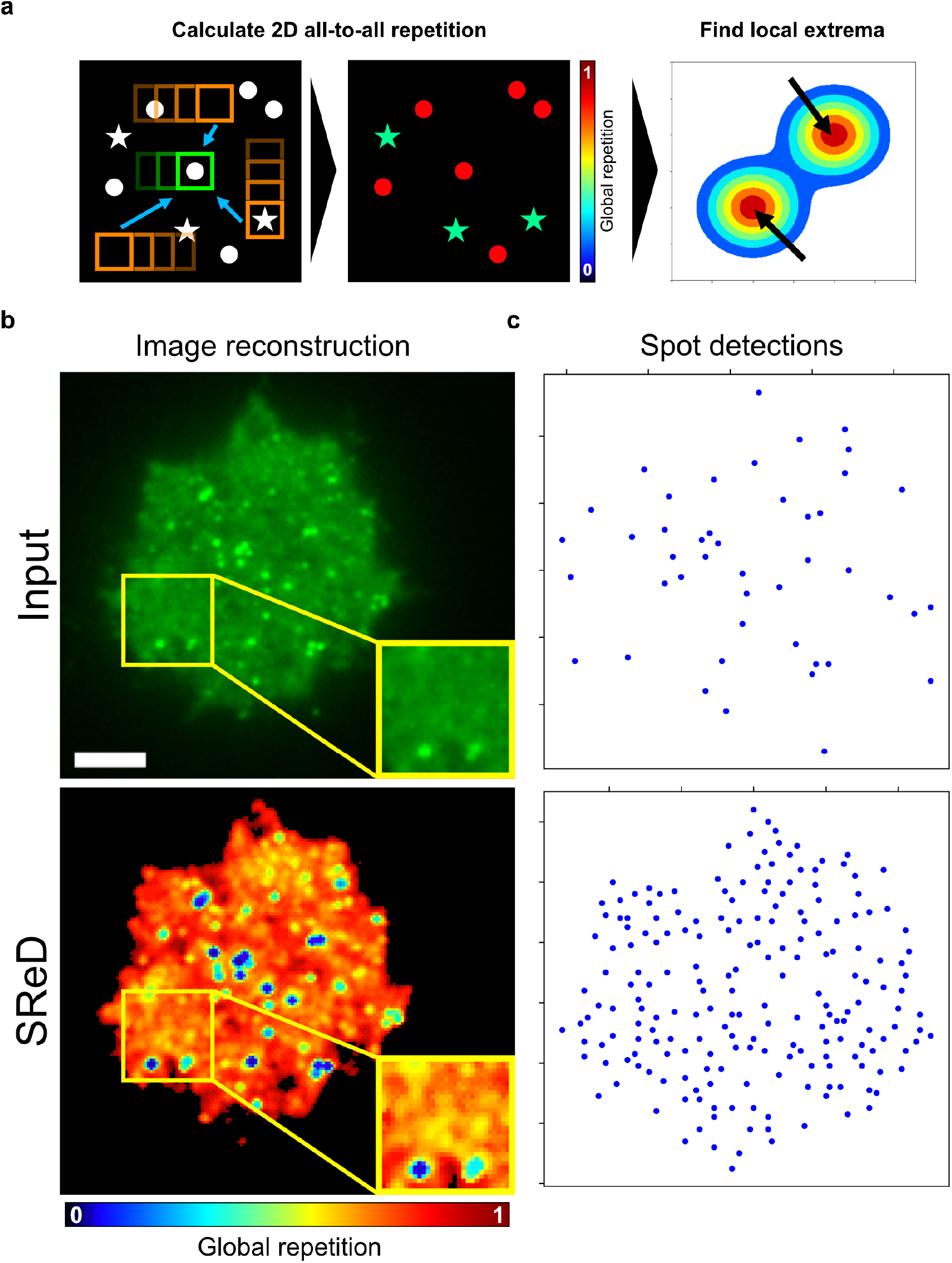
Detection of HIV-1 virus-like particles using Global Repetition. **a**, Workflow diagram illustrating the “all-to-all” sampling scheme in Global Repetition. The global repetition map is a platform for quantitative analysis such as local extrema calculation. **b**, Top panel: Image reconstruction of an acivated Jurkat cell expressing an HIV-1 Gag-EGFP construct, which induces the assembly of virus-like particles (VLPs). Scale bar: 5 µm. Bottom panel: Global repetition map calculated from the input image in the top panel, using a block size of 7×7 pixels and the modified cosine similarity metric. The insets in both panels highlight a region where assembling viral structures can be discerned in the global repetition map but not in the input image. **c**, Local maxima calculated from the input image and the (inverted) global repetition map, showing a higher number of detections in the global repetition map using the same prominence threshold.

**Fig. S5.**
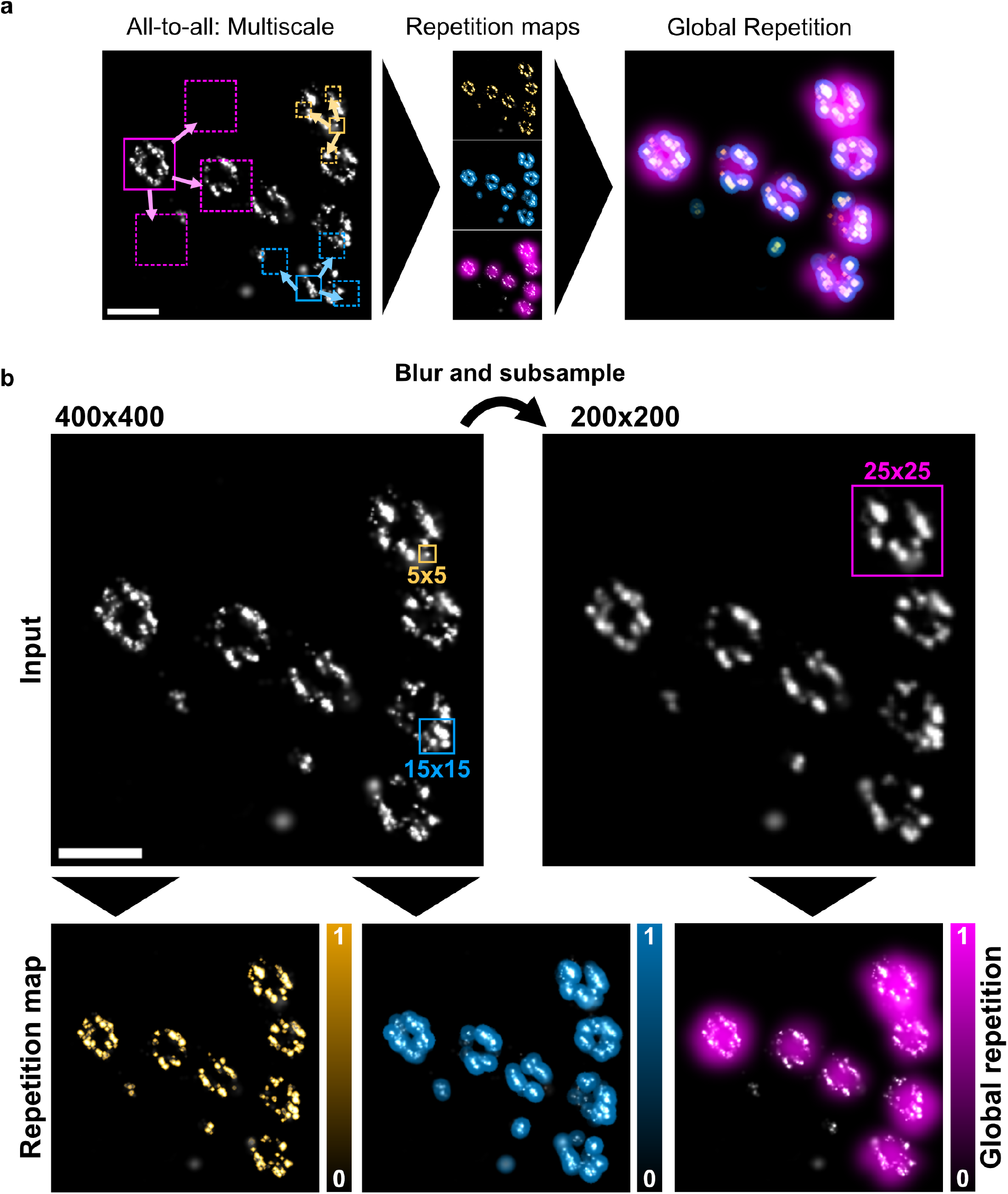
Multiscale detection of Nuclear Pore Complex structures using Global Repetition. **a**, Workflow diagram illustrating the modulation of block-to-image size ratio modulation for multiscale global repetition analysis. **b**, Image reconstructions of the input image and the block sizes used for global repetition analysis. A low- and intermediate-order analysis used the original image dimensions (400×400 pixels) and 5×5 and 15×15 pixel block sizes to detect single nucleoporins (orange) and nucleoporin clusters (blue). A high-order analysis used a downscaled input image (200×200 pixels) and a block size of 25×25 pixels to detect entire NPC units.

**Fig. S6.**
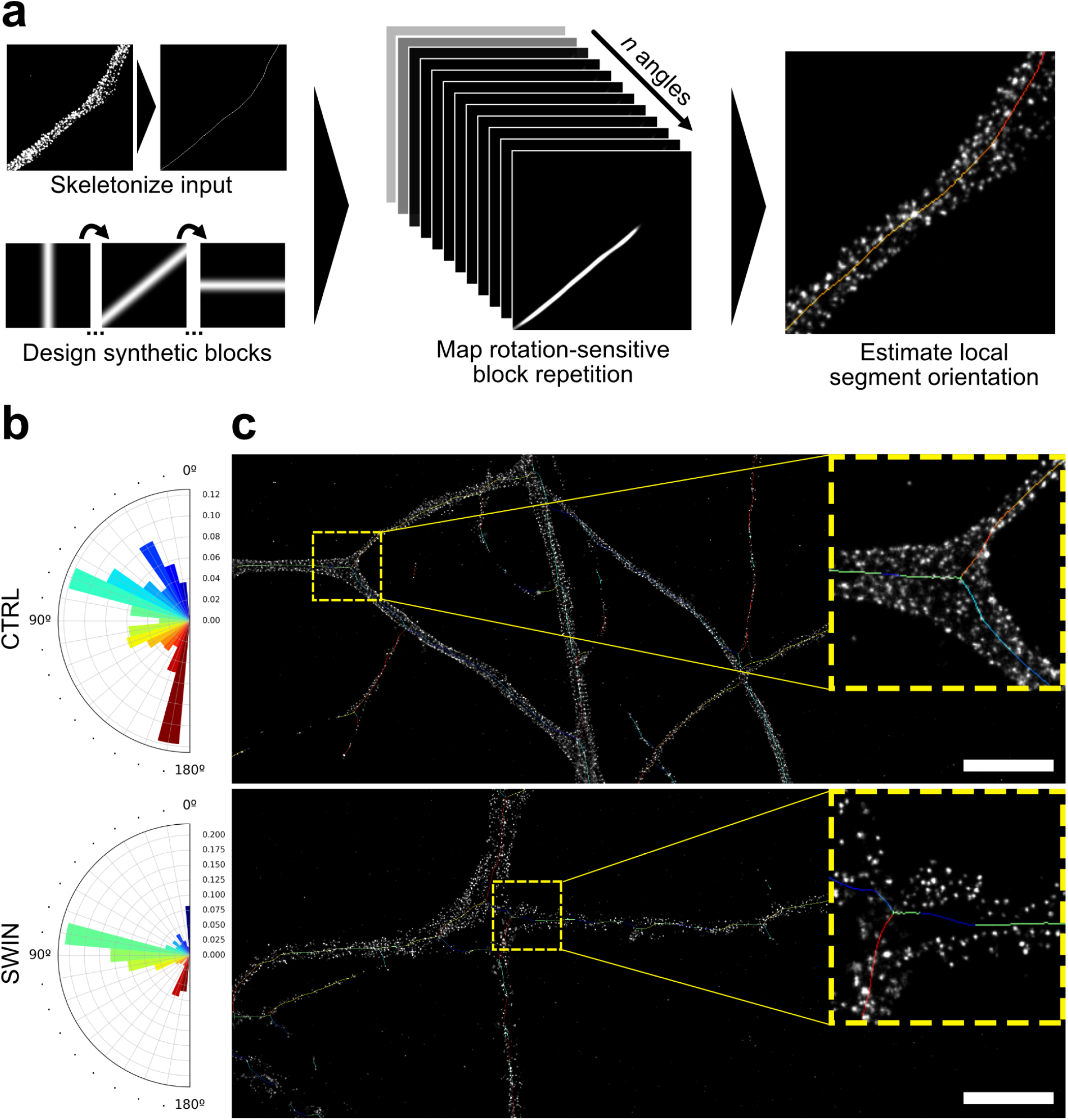
Automated estimation of axon orientations. **a**, Workflow diagram illustrating the estimation of axon orientations. Synthetic blocks comprising lines at different orientations are used to calculate repetitions in skeletonised axons. Each skeleton coordinate is labelled with the angle corresponding to the highest SRS. **b**, Polar plots depicting the distribution of axon angles in the control (top panel) and swinholide A (bottom panel) samples. **c**, Overlay of representative input images and their angle-labelled skeletons, using the same colour code used in panel b). Scale bar: 5 µm.

**Fig. S7.**
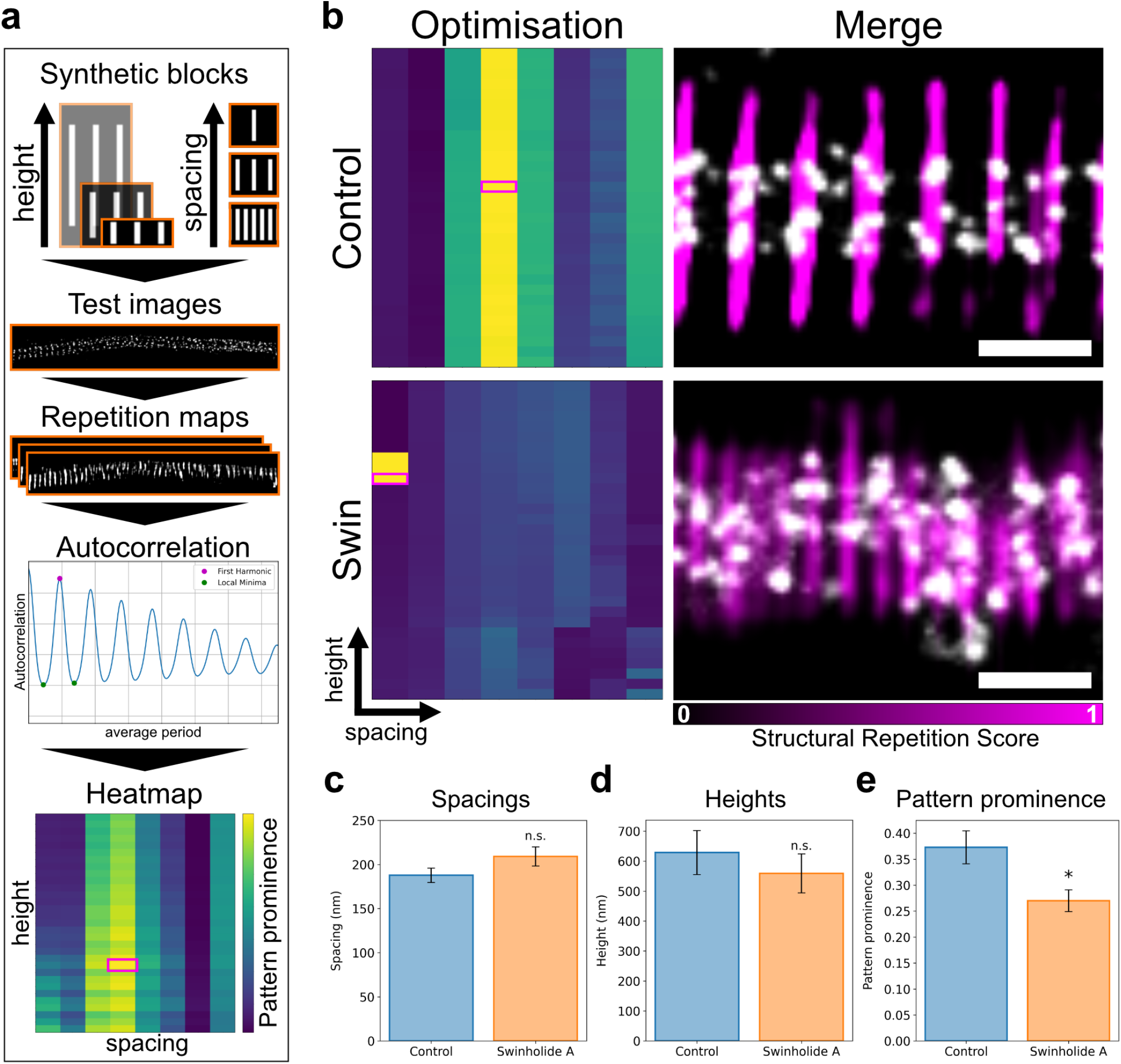
Optimisation of block parameters for ring pattern detection. **a**, Schematic representation of the tasks performed. A set of 248 synthetic blocks with different spacing and height combinations were generated. Test images (30 for each group) were randomly extracted from each dataset and SReD was used to calculate the repetition maps of each reference block. The repetition maps were analysed using autocorrelation functions and the patterns’ prominence (given by the autocorrelations’ first harmonic amplitude) was used to assess how well each reference block fitted the data. For each test image, this information is depicted in a heatmap. The combination of parameters with the highest pattern prominence value is chosen in from each heatmap and averaged within each dataset to calculate the optimised block. **b**, Representative examples of the optimisation process. The heatmaps where the highest pattern prominences were found are shown. Scale bar: 0.4 µm. **c-e**, Plots of the optimised parameters calculated by comparing the distributions of each parameter between groups (n=248, n.s. p>0.05, * p<0.05, Mann-Whitney U test).

**Fig. S8.**
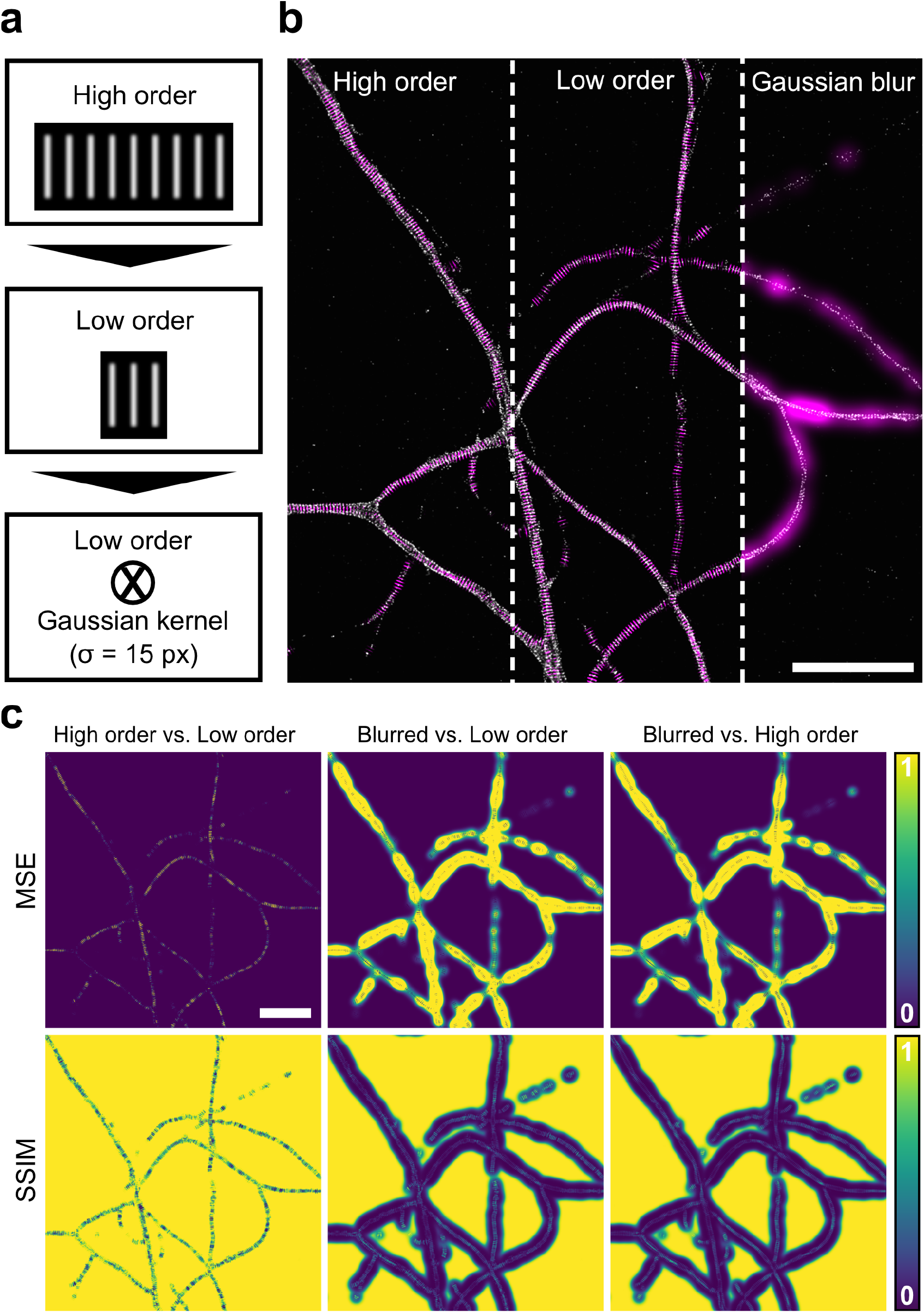
Detection of high-order ring patterns. **a**, Schematic representation of the samples generated. Top panel: Reference block containing a high-order ring pattern with 9 rings. Center panel: Reference block containing a low-order ring pattern with 3 rings. Bottom panel: Convolution with a Gaussian kernel (15 px radius). **b**, Overlay of a representative example of the input data and the corresponding block repetition maps. **c**, Error maps demonstrating that SReD enables detecting high-order patterns without discarding low-order information. The high-order repetition maps are a more robust representation of the low-order structures compared to the method using convolutions with Gaussian kernels. Scale bars: 5 µm.

**Fig. S9.**
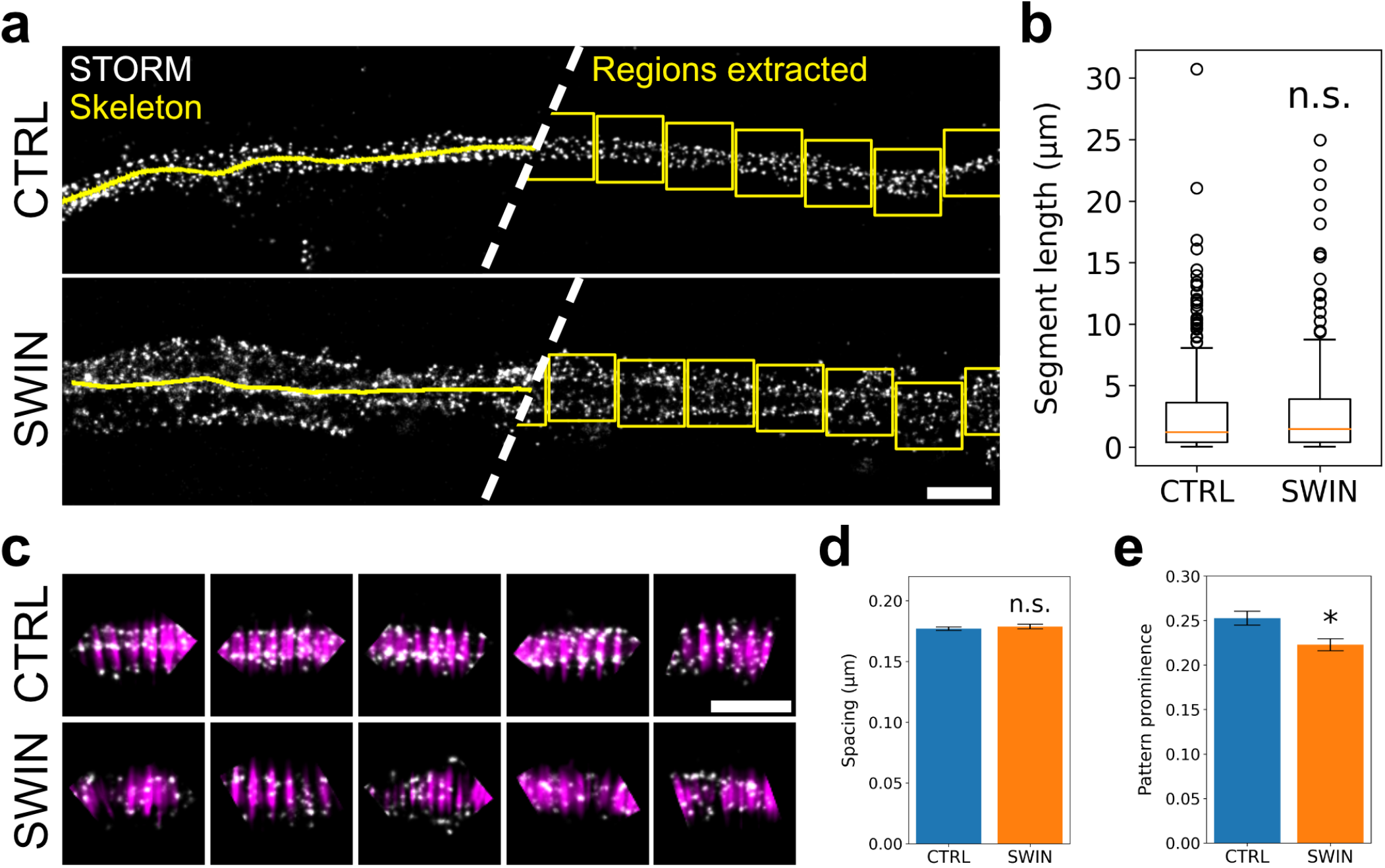
Quantitative analysis of ring patterns in axon segments. **a**, Image reconstructions illustrating the approach used to automatically extract non-overlapping regions along the axons’ skeletons. Scale bar: 1 µm. **b**, Length distributions of the skeletonised axon segments analysed. **c**, Representative examples of the non-overlapping regions extracted along the axons’ skeleton segments (STORM in gray, SReD repetition maps in magenta). Scale bar: 1 µm. **d**, Average spacing of the regions analysed (N=6, mean ± SEM - CTRL: 177 nm ± 1 nm, SWIN: 179 nm ± 2 nm, n.s. p>0.05, t-test). **e**, Pattern prominence of the regions analysed (N=6, mean ± SEM - CTRL: 0.253 ± 0.008, SWIN: 0.223 ± 0.007, * p<0.05, t-test).

**Fig. S10.**
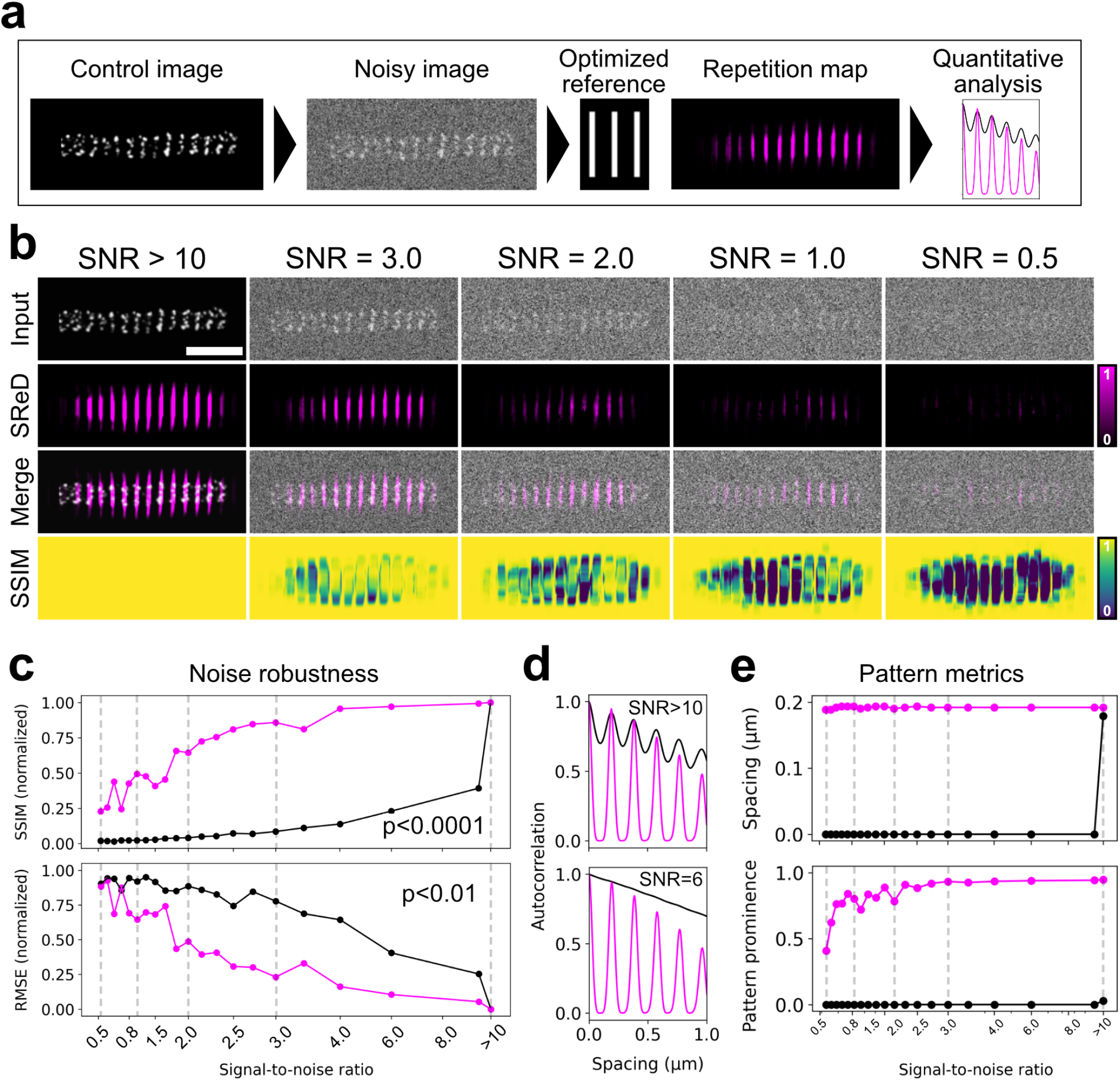
Evaluation of SReD’s performance in images containing random noise. **a**, Schematic of the tasks performed. A control image containing a periodic pattern is corrupted with random noise. The periodic pattern is mapped using an optimised reference block. The repetition maps are quantitatively analysed using autocorrelation functions. **b**, Visual comparison of image reconstructions at different signal-to-noise ratios (SNRs). Top row: Input images with SNRs decreasing from left to right. Middle row: Repetition maps calculated using an optimised reference block. Bottom row: Merge of the input images and the repetition maps. Scale bar: 1 µm **c**, Evaluation of noise robustness by comparing the distributions of control vs. noisy images (black) and the corresponding repetition maps (magenta) at different SNRs using different metrics. The repetition maps provided a superior representation of the control conditions in all cases. Top panel: Distributions calculated using the Structural Similarity Index (SSIM) metric (p<0.0001, t-test). Bottom panel: Distributions calculated using the Root Mean Squared Error (RMSE) metric (p<0.01, t-test). **d**, Autocorrelation plots demonstrating the sensitivity to the periodic patterns in the input images (black) and the SReD repetition maps (magenta). **e**, Distribution of the inter-ring spacing (top panel) and pattern prominence (bottom panel) calculated from the autocorrelation functions of the input data (black) and the SReD repetition maps (magenta) at different SNRs.

**Fig. S11.**
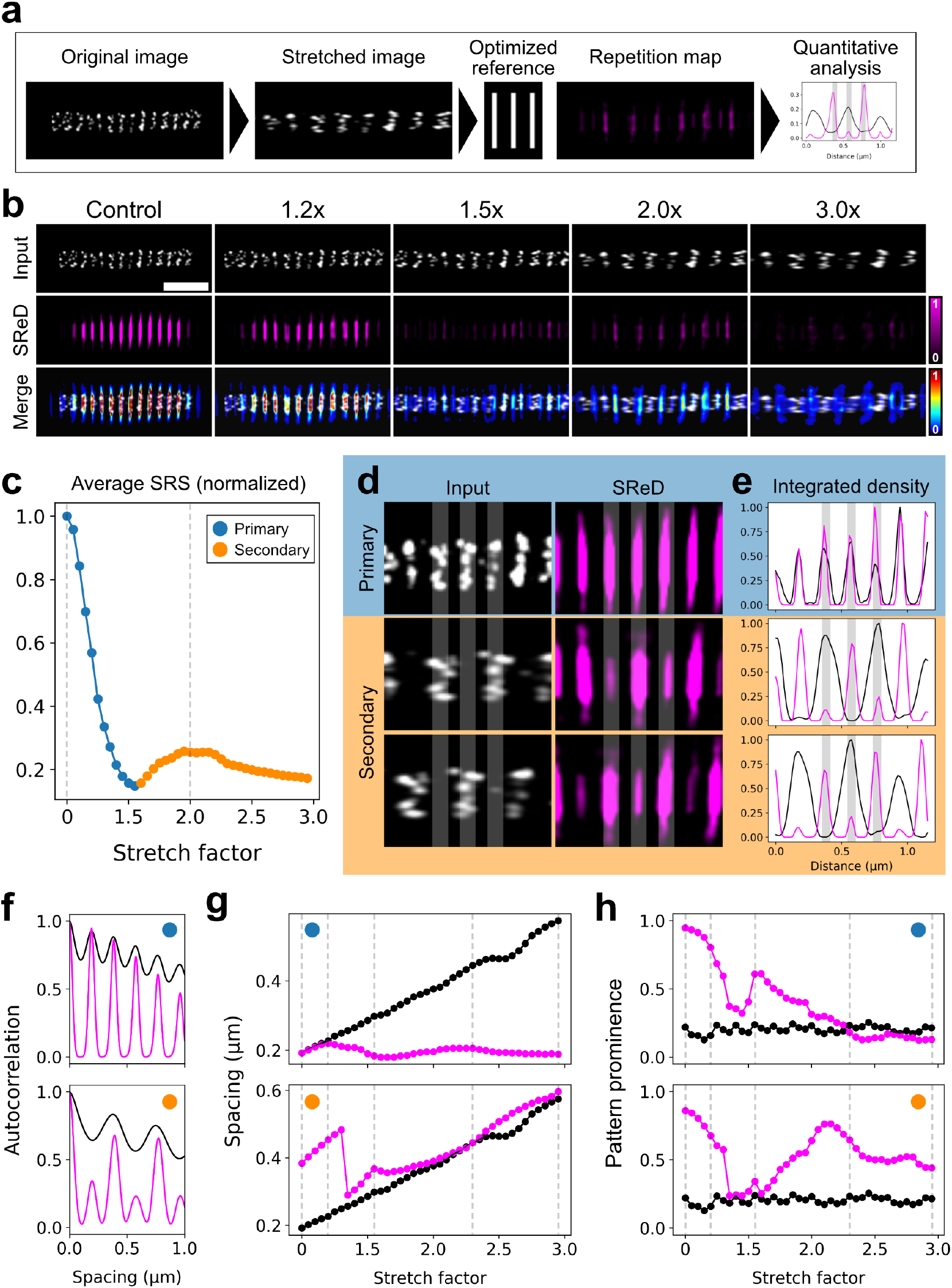
Evaluation of SReD’s specificity. **a**, Schematic of the tasks performed. A control image containing a periodic pattern is stretched along its width to disrupt the pattern’s characteristics. The periodic pattern is mapped using an optimised reference block. The repetition maps are quantitatively analysed using autocorrelation functions. **b**, Visual comparison of image reconstructions at different stretch factors. Top row: Input images with stretch factors increasing from left to right. Middle row: Repetition maps calculated using an optimised reference block. Bottom row: Merge of the input images and the repetition maps. Scale bar: 1 µm. **c**, Average Structural Repetition Factor (SRS) in the repetition maps plotted against the stretch factor. A primary pattern (blue) and a secondary pattern (orange) are detected. **d**, Image reconstructions of the primary and secondary patterns detected in c). **e**, 2D intensity profiles (i.e., integrated density) of the images in d). **f**, Autocorrelation plots of the input images (black) and repetition maps (magenta) at stretch factors 0 (top) and 2.0 (bottom). **g**, Inter-ring spacings calculated from the autocorrelation analysis plotted against the stretch factor. **h**, Pattern prominences calculated from the autocorrelation analysis plotted against the stretch factor. The dashed gray lines indicate the samples shown in panel b).

**Fig. S12.**
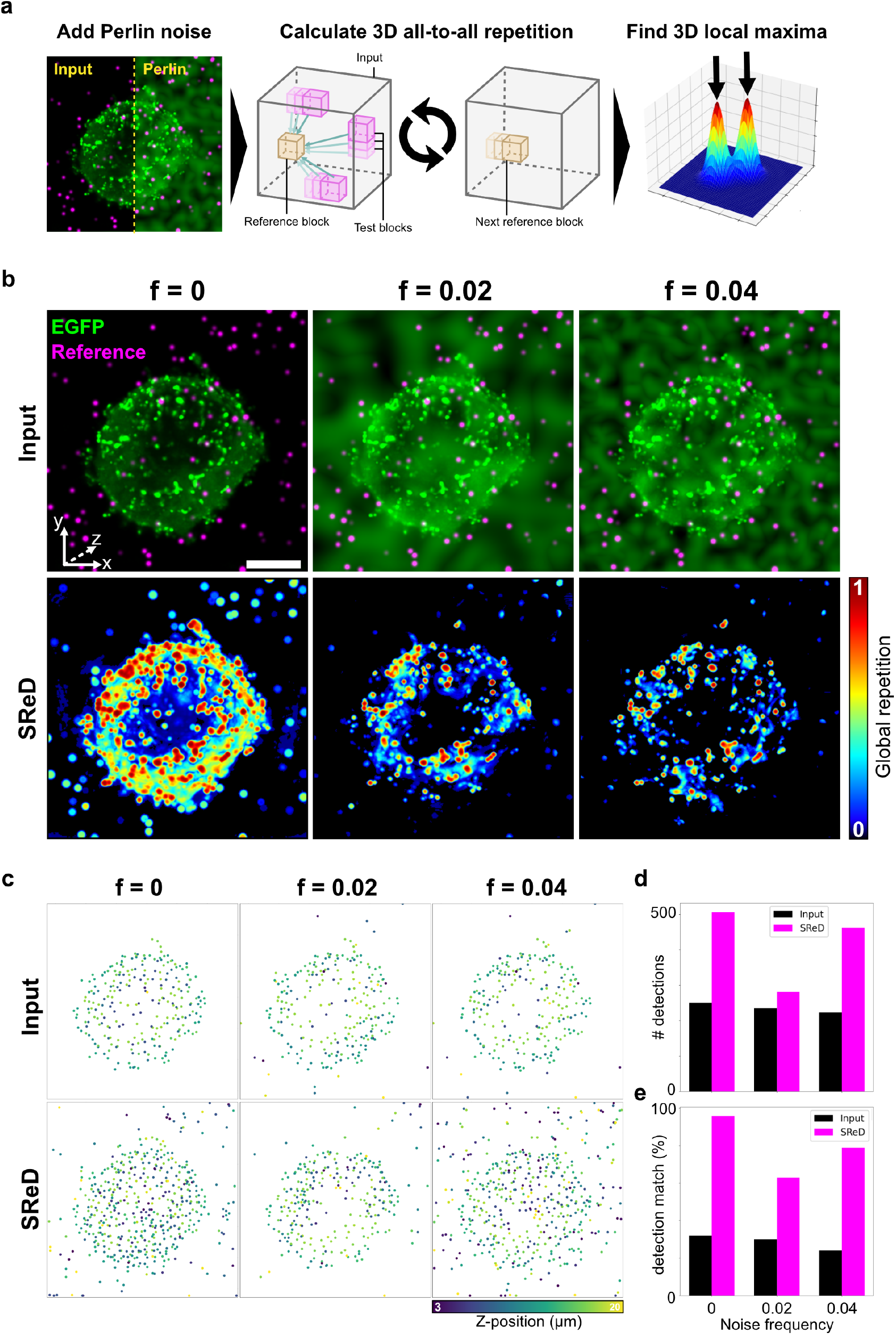
Evaluation of SReD’s performance in images corrupted with non-specific structures. **a**, Schematic of the tasks performed. Perlin noise is added to a 3D image containing a Jurkat cell expressing an HIV-1 Gag-EGFP construct (green) and synthetic bead structures resembling free viral particles (magenta). A Global Repetition map is calculated and 3D local maxima are calculated. **b**, Input images with Perlin noise of different frequencies and their corresponding Global Repetition maps. **c**, Plots of the spot detections obtained by calculating local maxima in the input images and their Global Repetition maps. **d**, Plot showing the number of detections obtained in each sample. **e**, Plot showing the percentage of reference detections obtained in each sample.

**Fig. S13.**
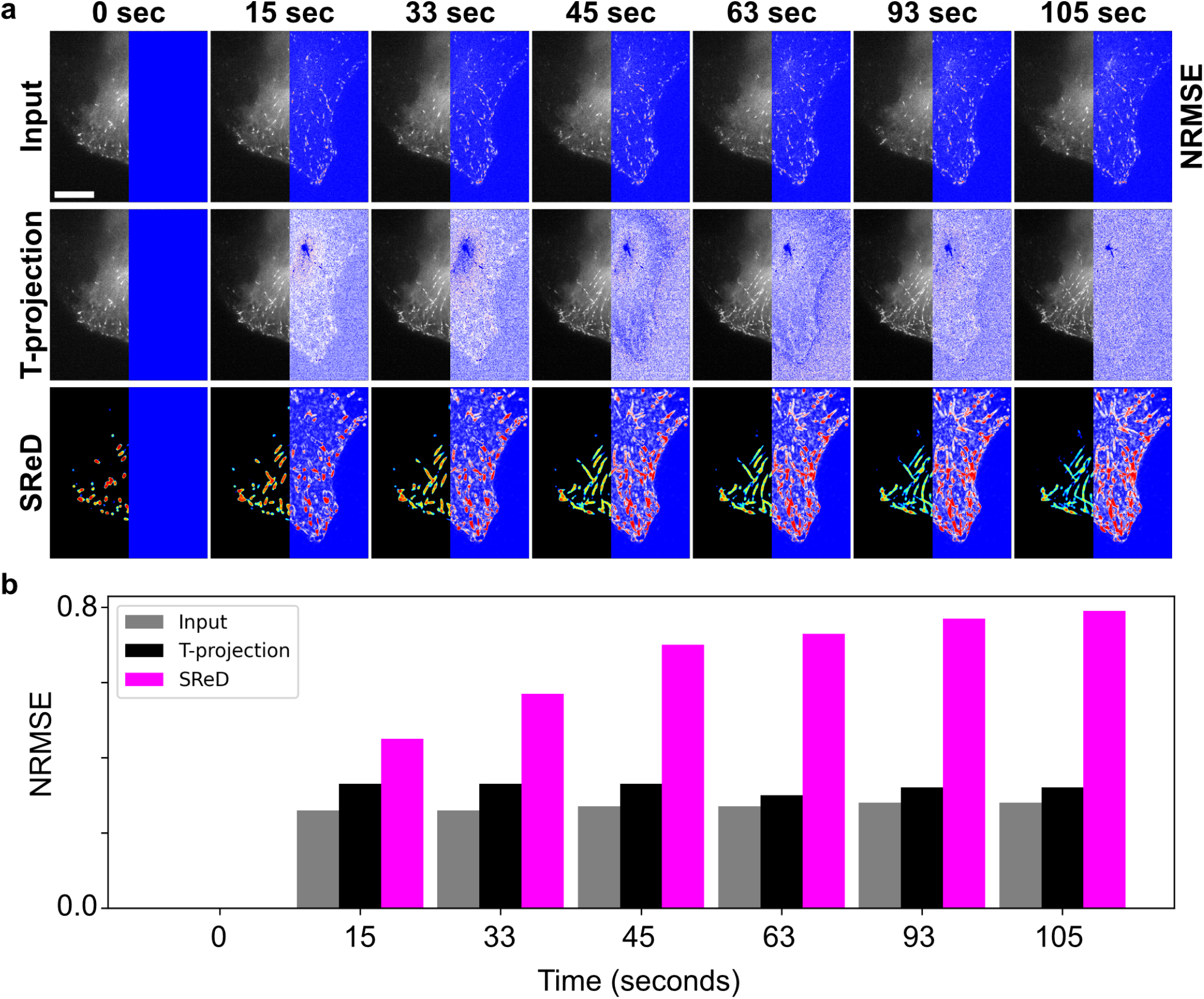
Assessment of microtubule dynamics using Global Repetition. **a**, Analysis of structural stability by comparing different time frames with the initial state (0 sec) using the Normalised Root Mean Squared Error (NRMSE) metric. Top row: Input images showing the final state in each time frame. Middle row: Time projections of all time points within each time frame. Bottom row: SReD global repetition maps calculated using time as the third dimension of the analysis. **b**, Plot showing the NRMSE distributions of each dataset.

## Notes

https://github.com/henriqueslab/sred

### Competing Interest Statement

The authors have declared no competing interest.

## Bibliography

1. Nam Hyeong Kim, Hojae Choi, Zafar Muhammad Shahzad, Heesoo Ki, Jaekyoung Lee, Heeyeop Chae, and Yong Ho Kim. Supramolecular assembly of protein building blocks: from folding to function. Nano Convergence, 9(1):4, January 2022. ISSN 2196-5404. doi: 10.1186/s40580-021-00294-3.

2. Afonso Mendes, Hannah S. Heil, Simao Coelho, Christophe Leterrier, and Ricardo Henriques. Mapping molecular complexes with super-resolution microscopy and single-particle analysis. Open Biology, 12(7):220079, July 2022. doi: 10.1098/rsob.220079. Publisher: Royal Society.

3. Anusha Aswath, Ahmad Alsahaf, Ben N. G. Giepmans, and George Azzopardi. Segmentation in large-scale cellular electron microscopy with deep learning: A literature survey. Medical Image Analysis, 89:102920, October 2023. ISSN 1361-8415. doi: 10.1016/j.media.2023.102920.

4. Olaf Ronneberger, Philipp Fischer, and Thomas Brox. U-Net: Convolutional Networks for Biomedical Image Segmentation, May 2015. arXiv:1505.04597 [cs].

5. Thorsten Falk, Dominic Mai, Robert Bensch, Özgün Çiçek, Ahmed Abdulkadir, Yassine Marrakchi, Anton Böhm, Jan Deubner, Zoe Jäckel, Katharina Seiwald, Alexander Dovzhenko, Olaf Tietz, Cristina Dal Bosco, Sean Walsh, Deniz Saltukoglu, Tuan Leng Tay, Marco Prinz, Klaus Palme, Matias Simons, Ilka Diester, Thomas Brox, and Olaf Ronneberger. U-Net: deep learning for cell counting, detection, and morphometry. Nature Methods, 16(1):67– 70, January 2019. ISSN 1548-7105. doi: 10.1038/s41592-018-0261-2. Publisher: Nature Publishing Group.

6. Romain F. Laine, Ignacio Arganda-Carreras, Ricardo Henriques, and Guillaume Jacquemet. Avoiding a replication crisis in deep-learning-based bioimage analysis. Nature Methods, 18(10):1136–1144, October 2021. ISSN 1548-7105. doi: 10.1038/s41592-021-01284-3. Publisher: Nature Publishing Group.

7. Hamidreza Heydarian, Florian Schueder, Maximilian T. Strauss, Ben van Werkhoven, Mohamadreza Fazel, Keith A. Lidke, Ralf Jungmann, Sjoerd Stallinga, and Bernd Rieger. Template-free 2D particle fusion in localization microscopy. Nature Methods, 15(10):781– 784, October 2018. ISSN 1548-7105. doi: 10.1038/s41592-018-0136-6. Publisher: Nature Publishing Group.

8. Hamidreza Heydarian, Maarten Joosten, Adrian Przybylski, Florian Schueder, Ralf Jungmann, Ben van Werkhoven, Jan Keller-Findeisen, Jonas Ries, Sjoerd Stallinga, Mark Bates, and Bernd Rieger. 3D particle averaging and detection of macromolecular symmetry in localization microscopy. Nature Communications, 12(1):2847, May 2021. ISSN 2041-1723. doi: 10.1038/s41467-021-22006-5. Publisher: Nature Publishing Group.

9. Jérôme Boulanger, Charles Kervrann, Patrick Bouthemy, Peter Elbau, Jean-Baptiste Sibarita, and Jean Salamero. Patch-Based Nonlocal Functional for Denoising Fluorescence Microscopy Image Sequences. IEEE Transactions on Medical Imaging, 29(2):442–454, February 2010. ISSN 1558-254X. doi: 10.1109/TMI.2009.2033991. Conference Name: IEEE Transactions on Medical Imaging.

10. Angélique Jimenez, Karoline Friedl, and Christophe Leterrier. About samples, giving examples: Optimized Single Molecule Localization Microscopy. Methods, 174:100–114, March 2020. ISSN 1046-2023. doi: 10.1016/j.ymeth.2019.05.008.

11. Olivier Burri and Romain Guiet. DAPI and Phase Contrast Images Dataset, May 2019.

12. Robin Diekmann, Maurice Kahnwald, Andreas Schoenit, Joran Deschamps, Ulf Matti, and Jonas Ries. Optimizing imaging speed and excitation intensity for single-molecule localization microscopy. Nature Methods, 17(9):909–912, September 2020. ISSN 1548-7105. doi: 10.1038/s41592-020-0918-5. Publisher: Nature Publishing Group.

13. Christophe Leterrier, Pankaj Dubey, and Subhojit Roy. The nano-architecture of the axonal cytoskeleton. Nature Reviews Neuroscience, 18(12):713–726, December 2017. ISSN 1471-0048. doi: 10.1038/nrn.2017.129. Publisher: Nature Publishing Group.

14. Christophe Leterrier. Putting the axonal periodic scaffold in order. Current Opinion in Neurobiology, 69:33–40, August 2021. ISSN 0959-4388. doi: 10.1016/j.conb.2020.12.015.

15. Ke Xu, Guisheng Zhong, and Xiaowei Zhuang. Actin, Spectrin, and Associated Proteins Form a Periodic Cytoskeletal Structure in Axons. Science, 339(6118):452–456, January 2013. doi: 10.1126/science.1232251. Publisher: American Association for the Advancement of Science.

16. Stéphane Vassilopoulos, Solène Gibaud, Angélique Jimenez, Ghislaine Caillol, and Christophe Leterrier. Ultrastructure of the axonal periodic scaffold reveals a braid-like organization of actin rings. Nature Communications, 10(1):5803, December 2019. ISSN 2041-1723. doi: 10.1038/s41467-019-13835-6. Publisher: Nature Publishing Group.

17. Sepiso K. Masenga, Bislom C. Mweene, Emmanuel Luwaya, Lweendo Muchaili, Makondo Chona, and Annet Kirabo. HIV–Host Cell Interactions. Cells, 12(10):1351, January 2023. ISSN 2073-4409. doi: 10.3390/cells12101351. Number: 1. Publisher: Multidisciplinary Digital Publishing Institute.

18. Wolfgang Hübner, Ping Chen, Armando Del Portillo, Yuxin Liu, Ronald E. Gordon, and Benjamin K. Chen. Sequence of Human Immunodeficiency Virus Type 1 (HIV-1) Gag Localization and Oligomerization Monitored with Live Confocal Imaging of a Replication-Competent, Fluorescently Tagged HIV-1. Journal of Virology, 81(22):12596–12607, November 2007. ISSN 0022-538X. doi: 10.1128/JVI.01088-07.

19. Charlotte Floderer, Jean-Baptiste Masson, Elise Boilley, Sonia Georgeault, Peggy Merida, Mohamed El Beheiry, Maxime Dahan, Philippe Roingeard, Jean-Baptiste Sibarita, Cyril Favard, and Delphine Muriaux. Single molecule localisation microscopy reveals how HIV-1 Gag proteins sense membrane virus assembly sites in living host CD4 T cells. Scientific Reports, 8:16283, November 2018. ISSN 2045-2322. doi: 10.1038/s41598-018-34536-y.

20. Tatiana Stepanova, Jenny Slemmer, Casper C. Hoogenraad, Gideon Lansbergen, Bjorn Dortland, Chris I. De Zeeuw, Frank Grosveld, Gert van Cappellen, Anna Akhmanova, and Niels Galjart. Visualization of Microtubule Growth in Cultured Neurons via the Use of EB3-GFP (End-Binding Protein 3-Green Fluorescent Protein). Journal of Neuroscience, 23(7):2655–2664, April 2003. ISSN 0270-6474, 1529-2401. doi: 10.1523/JNEUROSCI.23-07-02655.2003. Publisher: Society for Neuroscience Section: ARTICLE.

21. Wenjiang Liu, Tao Liu, Mengtian Rong, Ruolin Wang, and Hao Zhang. A fast noise variance estimation algorithm. In 2011 Asia Pacific Conference on Postgraduate Research in Microelectronics & Electronics, pages 61–64, October 2011. doi: 10.1109/PrimeAsia.2011.6075071. ISSN: 2159-2160.

22. João I. Mamede, Joseph Griffin, Stéphanie Gambut, and Thomas J. Hope. A New Generation of Functional Tagged Proteins for HIV Fluorescence Imaging. Viruses, 13(3):386, March 2021. ISSN 1999-4915. doi: 10.3390/v13030386. Number: 3. Publisher: Multidisciplinary Digital Publishing Institute.

23. William Hadley Richardson. Bayesian-Based Iterative Method of Image Restoration*. JOSA, 62(1):55–59, January 1972. doi: 10.1364/JOSA.62.000055. Publisher: Optica Publishing Group.

24. L. B. Lucy. An iterative technique for the rectification of observed distributions. The Astronomical Journal, 79:745, June 1974. ISSN 0004-6256. doi: 10.1086/111605. Publisher: IOP ADS Bibcode: 1974AJ 79.745L.

25. Stephen D. Carter, João I. Mamede, Thomas J. Hope, and Grant J. Jensen. Correlated cryogenic fluorescence microscopy and electron cryo-tomography shows that exogenous TRIM5α can form hexagonal lattices or autophagy aggregates in vivo. Proceedings of the National Academy of Sciences, 117(47):29702–29711, November 2020. doi: 10.1073/pnas.1920323117. Publisher: Proceedings of the National Academy of Sciences.

26. Sunnie M. Yoh, João I. Mamede, Derrick Lau, Narae Ahn, Maria T. Sánchez-Aparicio, Joshua Temple, Andrew Tuckwell, Nina V. Fuchs, Gianguido C. Cianci, Laura Riva, Heather Curry, Xin Yin, Stéphanie Gambut, Lacy M. Simons, Judd F. Hultquist, Renate König, Yong Xiong, Adolfo García-Sastre, Till Böcking, Thomas J. Hope, and Sumit K. Chanda. Recognition of HIV-1 capsid by PQBP1 licenses an innate immune sensing of nascent HIV-1 DNA. Molecular Cell, 82(15):2871–2884.e6, August 2022. ISSN 10972765. doi: 10.1016/j.molcel.2022.06.010.

27. Nobuyuki Otsu. A Threshold Selection Method from Gray-Level Histograms. IEEE Transactions on Systems, Man, and Cybernetics, 9(1):62–66, January 1979. ISSN 2168-2909. doi: 10.1109/TSMC.1979.4310076. Conference Name: IEEE Transactions on Systems, Man, and Cybernetics.

28. E. Meijering, M. Jacob, J.-C.f. Sarria, P. Steiner, H. Hirling, and M. Unser. Design and validation of a tool for neurite tracing and analysis in fluorescence microscopy images. Cytometry Part A, 58A(2):167–176, 2004. ISSN 1552-4930. doi: 10.1002/cyto.a.20022. _eprint: https://onlinelibrary.wiley.com/doi/pdf/10.1002/cyto.a.20022.

## Supplementary Bibliography

1. Jérôme Boulanger, Charles Kervrann, Patrick Bouthemy, Peter Elbau, Jean-Baptiste Sibarita, and Jean Salamero. Patch-Based Nonlocal Functional for Denoising Fluorescence Microscopy Image Sequences. IEEE Transactions on Medical Imaging, 29(2):442–454, February 2010. ISSN 1558-254X. doi: 10.1109/TMI.2009.2033991. Conference Name: IEEE Transactions on Medical Imaging.

2. Wenjiang Liu, Tao Liu, Mengtian Rong, Ruolin Wang, and Hao Zhang. A fast noise variance estimation algorithm. In 2011 Asia Pacific Conference on Postgraduate Research in Microelectronics & Electronics, pages 61–64, October 2011. doi: 10.1109/PrimeAsia.2011.6075071. ISSN: 2159-2160.

3. Angélique Jimenez, Karoline Friedl, and Christophe Leterrier. About samples, giving examples: Optimized Single Molecule Localization Microscopy. Methods, 174:100–114, March 2020. ISSN 1046-2023. doi: 10.1016/j.ymeth.2019.05.008.

4. Olivier Burri and Romain Guiet. DAPI and Phase Contrast Images Dataset, May 2019.

5. Nobuyuki Otsu. A Threshold Selection Method from Gray-Level Histograms. IEEE Transactions on Systems, Man, and Cybernetics, 9(1):62–66, January 1979. ISSN 2168-2909. doi: 10.1109/TSMC.1979.4310076. Conference Name: IEEE Transactions on Systems, Man, and Cybernetics.

6. E. Meijering, M. Jacob, J.-C.f. Sarria, P. Steiner, H. Hirling, and M. Unser. Design and validation of a tool for neurite tracing and analysis in fluorescence microscopy images. Cytometry Part A, 58A (2):167–176, 2004. ISSN 1552-4930. doi: 10.1002/cyto.a.20022. _eprint: https://onlinelibrary.wiley.com/doi/pdf/10.1002/cyto.a.20022.

7. Stéphane Vassilopoulos, Solène Gibaud, Angélique Jimenez, Ghislaine Caillol, and Christophe Leterrier. Ultrastructure of the axonal periodic scaffold reveals a braid-like organization of actin rings. Nature Communications, 10(1):5803, December 2019. ISSN 2041-1723. doi: 10.1038/s41467-019-13835-6. Publisher: Nature Publishing Group.

